# Recruitment of KRAS downstream target ARL4C to membrane protrusions accelerates pancreatic cancer cell invasion

**DOI:** 10.1101/2021.02.23.432112

**Authors:** Akikazu Harada, Shinji Matsumoto, Yoshiaki Yasumizu, Toshiyuki Akama, Hidetoshi Eguchi, Akira Kikuchi

## Abstract

Pancreatic cancer (PC) has a high mortality rate due to metastasis. Whereas KRAS is mutated in most PC patients, controlling KRAS or its downstream effectors has not been succeeded clinically. ARL4C is a small G protein whose expression is induced by the Wnt and EGF–RAS pathways. In the present study, we found that ARL4C is frequently overexpressed in PC patients and showed that its unique localization to membrane protrusions is required for cancer cell invasion. IQGAP1 was identified as a novel interacting protein for ARL4C. ARL4C recruited IQGAP1 and its downstream effector, MMP14, to membrane protrusions. Specific localization of ARL4C, IQGAP1, and MMP14 was the active site of invasion, which induced degradation of the extracellular matrix. Moreover, subcutaneously injected antisense oligonucleotide (ASO) against ARL4C into tumor-bearing mice suppressed metastasis of PC. These results suggest that ARL4C–IQGAP1–MMP14 signaling is activated at membrane protrusions of PC cells.

## Introduction

Pancreatic cancer is extremely aggressive and exhibits poor prognosis, with a 5-year survival of only 5%(Klein, 2013). Most pancreatic cancer-related deaths are due to metastatic disease, and more than 80% of patients have either locally advanced or metastatic disease(Hidalgo, 2010; Klein, 2013). Genome sequencing analysis has revealed the mutational landscape of pancreatic cancer and KRAS mutations are considered an initiating event in pancreatic ductal cells(Collins et al, 2012; Waddell et al, 2015). Irrespective of our improved understanding of tumor biology, the treatment outcome has not changed in the past 30 years. Therefore, new innovative treatment options need to be tested based on better understanding of the characteristics of pancreatic cancer.

ARL4C is a member of the ADP-ribosylation factor (ARF)-like protein (ARL) family, which belongs to the ARF protein subgroup of the small GTP-binding protein superfamily(Engel et al, 2004; Matsumoto et al, 2017; Wei et al, 2009). Cytohesin2/ARF nucleotide-binding site opener (ARNO), a GDP/GTP exchange factor of ARF family proteins, has been identified as a direct effector protein(Hofmann et al, 2007). ARL4C is expressed through activation of Wnt–β-catenin and EGF– RAS signaling and plays important roles in both epithelial morphogenesis and tumorigenesis(Matsumoto et al, 2017; Matsumoto et al, 2014). Because aberrant activation of the Wnt– β-catenin and/or EGF–RAS pathways are frequently observed in various types of cancers, ARL4C is indeed expressed in a number of cancers(Fujii et al, 2015; Fujii et al, 2016). In colon and lung cancer cells, ARL4C promotes cell proliferation through ARF6, RAC, RHO, and YAP/TAZ. On the other hand, in liver cancer cells, ARL4C promotes cell proliferation through phosphatidylinositol 3 kinase δ (PI3Kδ)(Harada et al, 2019). Thus, ARL4C would activate different downstream pathways in a cancer cell context-dependent manner. These prompted us to study the involvement of ARL4C, as a KRAS downstream molecule, in aggressiveness of pancreatic cancer, and IQGAP1 was identified as a binding protein of ARL4C.

IQ-domain GTPase-activation proteins (IQGAPs) are an evolutionally conserved family of proteins that bind to a diverse array of signaling and structural proteins(Hedman et al, 2015). Mammalian IQGAP1 is a well-characterized member of the IQGAP family and a fundamental regulator of cytoskeletal function(Briggs & Sacks, 2003). IQGAP1 is highly expressed in the tumor lesions and suggested to be involved in cancer cell metastasis(Johnson et al, 2009). (Sakurai-Yageta et al, 2008)Here, we show that ARL4C bound to IQGAP1 and recruited IQGAP1 and membrane type1-matrix metalloproteinase (MT1-MMP, also called MMP14)(Sakurai-Yageta et al, 2008) to membrane protrusions in a phosphatidylinositol (3,4,5)-trisphosphate (PIP3)-dependent manner and accelerated invasion. In addition, ARL4C antisense oligonucleotide (ASO) suppressed the lymph node metastases of pancreatic cancer cells orthotopically implanted into the pancreas of immunodeficient mice. These results suggest that the ARL4C–IQGAP1–MMP14 signaling axis promotes pancreatic cancer aggressiveness and that ARL4C is a novel molecular target for the treatment of pancreatic cancer.

## Results

### ARL4C is expressed in human pancreatic cancer

Whether ARL4C is expressed in pancreatic cancer patients was examined using immunohistochemistry. Fifty-seven pancreatic ductal adenocarcinoma (PDAC) patients without preoperative chemotherapy were classified into two groups, depending on ARL4C expression levels (high and low) (Figure 1A). High expression of ARL4C was observed in 47 cases (82%), but minimally detected in non-tumor regions of pancreatic ducts (Figure 1A). Anti-ARL4C antibody used in this study was validated in Western blotting and immunohistochemical assay (IHC) (Figure 1-figure supplement 1A and B). A significant difference was observed between low and high ARL4C expression based on perineural invasion (Supplementary file 1 Table 1). Because the perineural invasion is considered as one of the causes of the recurrence and metastasis after pancreatic resection (Liang et al, 2016), ARL4C expression may be correlated with the ability of cancer cell invasion.Consistently, ARL4C expression was correlated with decreased overall survival (Figure 1B). Similar results were obtained in the analysis of TCGA and GTEx datasets (Figure 1C and D). Univariate and multivariate analysis revealed that higher ARL4C expression was an independent prognostic factor (Table 1). Taken together, these results indicate that high expression of ARL4C is correlated with the aggressiveness and poor prognosis of pancreatic cancer.

**Figure 1.**
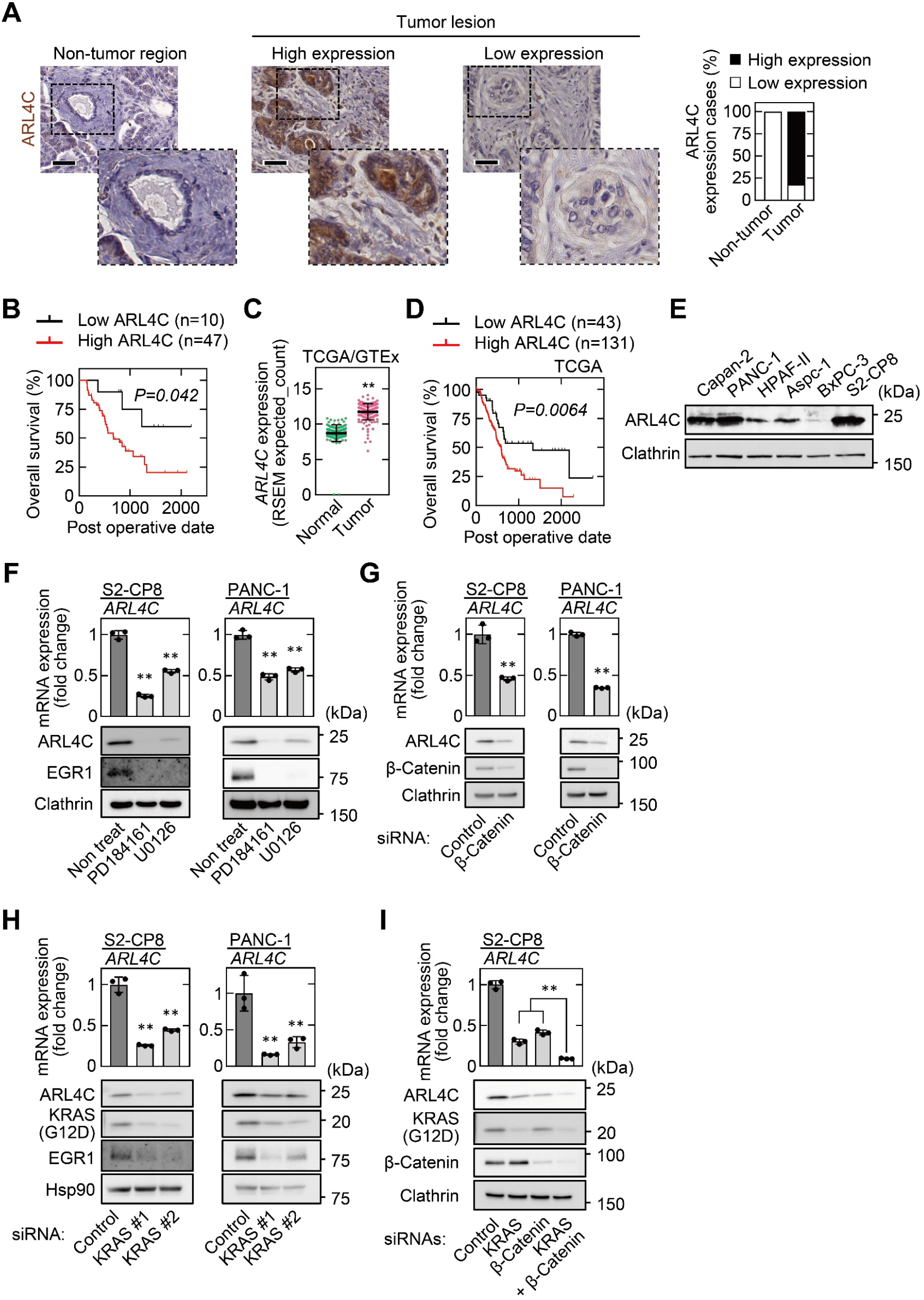
ARL4C is expressed in human pancreatic cancer. **A**, PDAC tissues (n = 57) were stained with anti-ARL4C antibody and hematoxylin. The percentages of ARL4C expression cases in the non-tumor regions and tumor lesions are shown. **B**, The relationship between overall survival and ARL4C expression in patients with PDAC. **C**, *ARL4C* mRNA levels in pancreatic adenocarcinoma and normal pancreatic tissues were analyzed using TCGA and GTEx datasets. The results shown are scatter plots with the mean ± s.e.m. *P* values were calculated using a two-tailed Student’s t-test. **D**, TCGA RNA sequencing and clinical outcome data for pancreatic cancer were analyzed. **E**, Lysates of the indicated pancreatic cancer cells were probed with the indicated antibodies. **F**, S2-CP8 and PANC-1 cells were treated with 10 μM PD184161 or 10 μM U0126, and *ARL4C* mRNA levels were measured by quantitative real-time PCR. Relative *ARL4C* mRNA levels were normalized to those of *GAPDH* and expressed as fold changes compared with the levels in control cells. Lysates were probed with the indicated antibodies. **G–I**, S2-CP8 cells and PANC-1 cells were transfected with the indicated siRNAs, and *ARL4C* mRNA levels were measured by quantitative real-time PCR. Relative *ARL4C* mRNA levels were normalized to those of *β2-microglobulin* and expressed as fold changes compared with the levels in control cells. Lysates were probed with the indicated antibodies. *EGR1* was used as an established transcription target gene of RAS signaling. **B,D,** Data were analyzed using Kaplan–Meier survival curves, and a log-rank test was used for statistical analysis. **F–I**, Data are shown as the mean ± s.d. of 3 independent experiments. *P* values were calculated using a two-tailed Student’s t-test (**G**) or one-way ANOVA followed by Bonferroni post hoc test (**F,H,I**). Scale bars in **A**, 50 μm. **, *P* < 0.01. See Figure 1-source data 1.

**Table 1.**
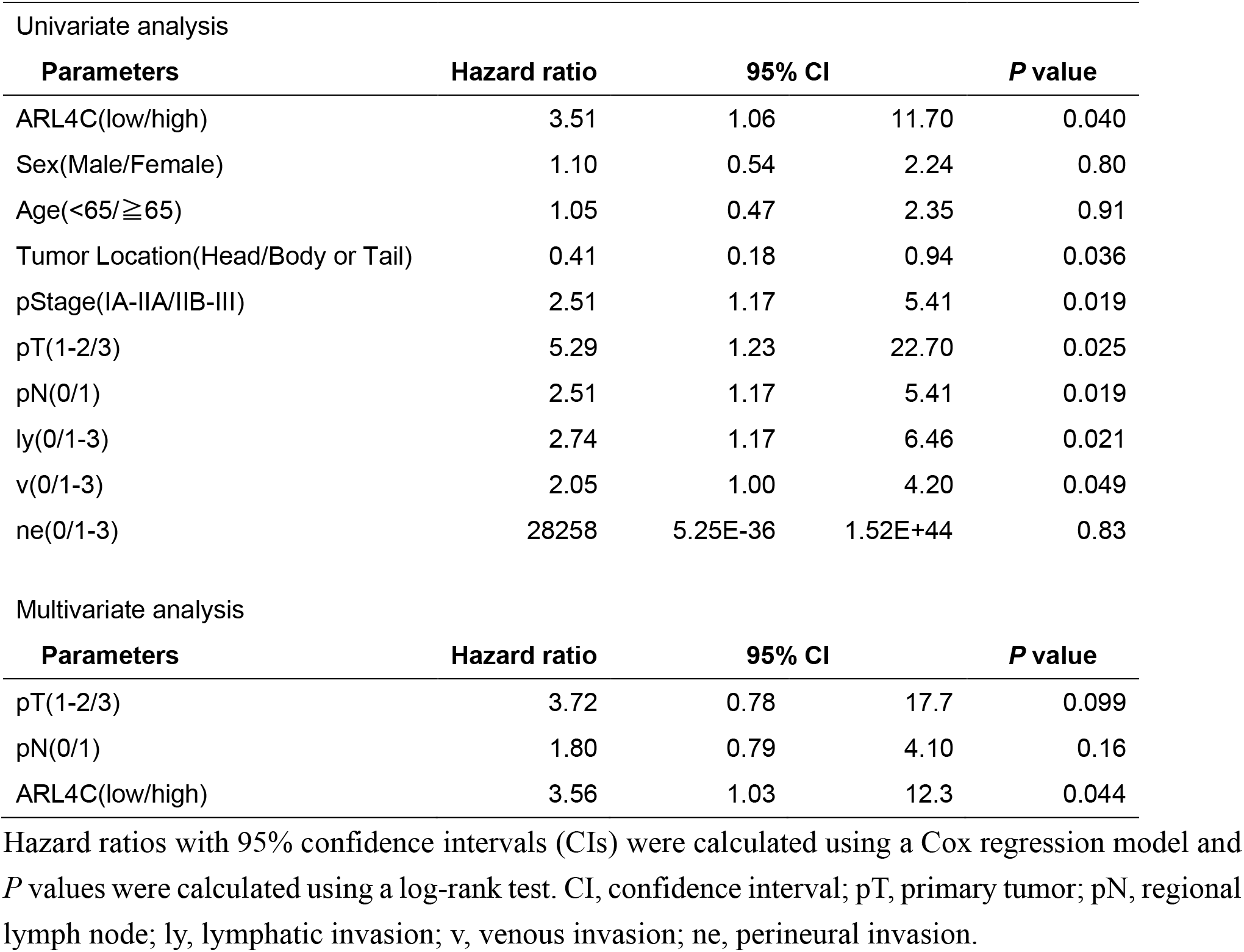
Univariate analysis and multivariate analysis of overall survival by Cox’s Proportional Hazard model.

In cultured pancreatic cancer cell lines, ARL4C was highly expressed in S2-CP8 and PANC-1 cells and it was barely detected in BxPC-3 cells (Figure 1E). Consistent with the previous results with IEC6 rat intestinal epithelial cells and colorectal and lung cancer cells(Fujii et al, 2015; Matsumoto et al, 2014), the MEK inhibitors PD184161 and U0126 and siRNAs for β-catenin and KRAS decreased ARL4C expression in S2-CP8 and PANC-1 cells (Figure 1F–H). In addition, simultaneous knockdown of KRAS and β-catenin further suppressed ARL4C expression (Figure 1I). Taken together, these results suggest that ARL4C is expressed in pancreatic cancer cells through activated RAS–MAP kinase and Wnt–β-catenin pathways.

### ARL4C expression is involved in the invasion of pancreatic cancer cells

ARL4C ASO-1316 has been shown to inhibit growth of xenograft tumors induced by colon and lung cancer cells(Harada et al, 2019; Kimura et al, 2020). However, ARL4C ASO-1316 had little effect on sphere formation of pancreatic cancer cell (Figure 2-figure supplement 1A). Since the clinicopathological analysis of human pancreatic cancer specimens indicates that ARL4C expression may be correlated with invasive ability, migratory and invasive abilities of S2-CP8 and PANC-1 cells were studied in Boyden chamber assays. ARL4C ASO-1316 inhibited the migratory and invasive abilities with dominant effects on invasion (Figure 2A and B; Figure 2-figure supplement 1B). Inhibition of migratory and invasive abilities by ARL4C ASO, targeting the non-coding region of *ARL4C* mRNA, was not observed in the cells expressing ARL4C-GFP ectopically (Figure 2C and D; Figure 2-figure supplement 1C). ARL4C is unique in that it is locked to the GTP-bound active form and localized to membrane protrusions of IEC6 and Madin-Darby canine kidney (MDCK) cells(Matsumoto et al, 2014). ARL4C-GFP was localized to membrane protrusions of S2-CP8 cells under Matrigel-coated 2D culture conditions (Figure 2-figure supplement 1D). ARL4C^Q72L^-GFP, in which the amino acid at the same position in a constitutively active RAS mutant was mutated, showed a similar distribution to ARL4C-GFP, but ARL4C^T27N^-GFP, which is an inactive form(Hofmann et al, 2007), did not (Figure 2-figure supplement 1D).

**Figure 2.**
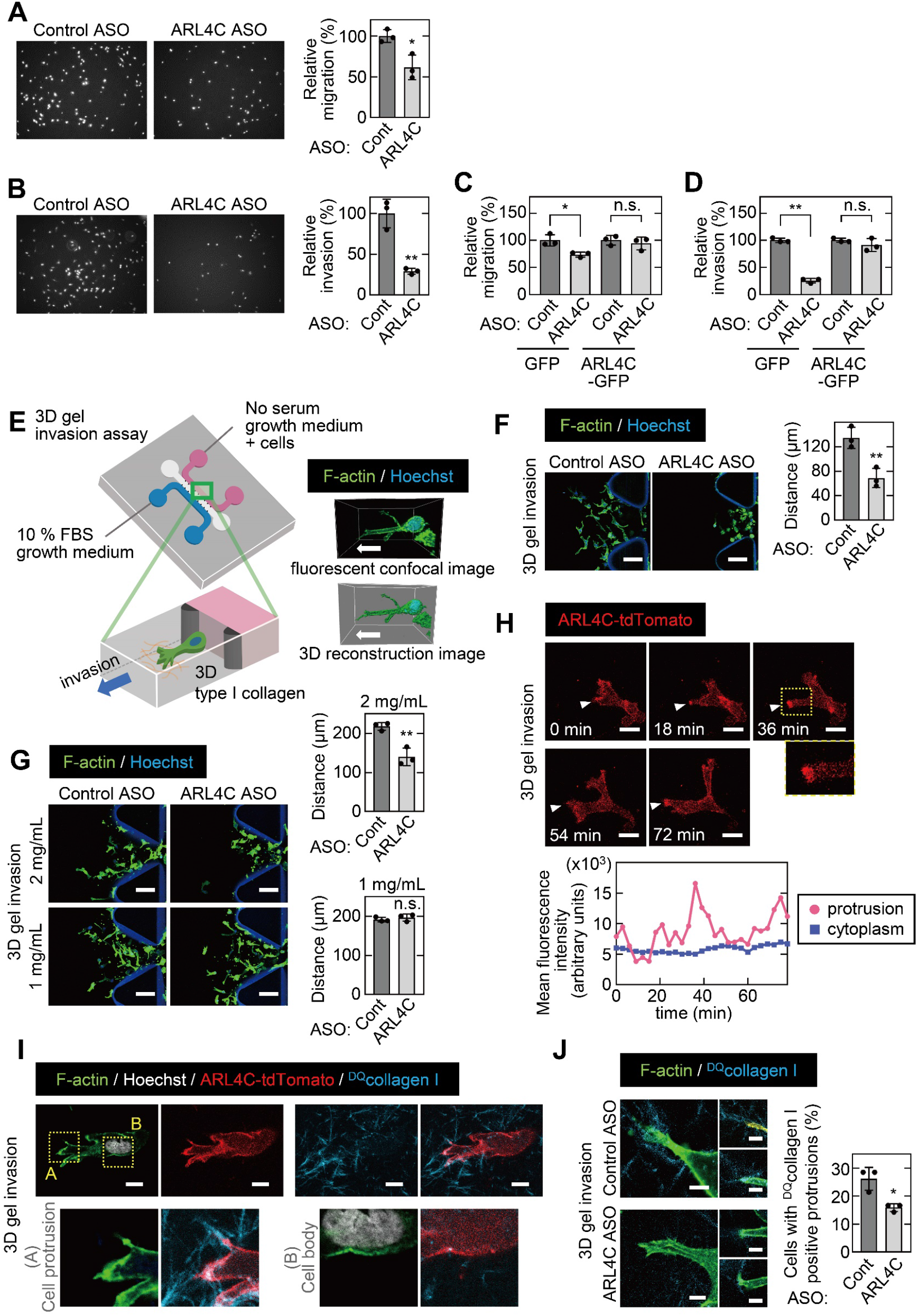
ARL4C expression is especially involved in the invasion of pancreatic cancer cells. **A-D,** S2-CP8 cells (**A,B**) or S2-CP8 cells expressing GFP or ARL4C-GFP (**C,D**) were transfected with control or ARL4C ASO-1316 and subjected to migration (**A,C**) and invasion (**B,D**) assays. Migratory and invasive abilities are expressed as the percentage of the same cells transfected with control ASO. **E**, A schematic illustration of 3D invasion into collagen I gel using a 3D cell culture chip is shown. There is a chemical concentration gradient across the gel channel and cells can invade into the gel. The right panel shows a fluorescent confocal image (top) and a 3D reconstructed image (bottom). **F**, S2-CP8 cells were transfected with control or ARL4C ASO-1316 and subjected to a 3D collagen I gel (2 mg/mL) invasion assay. The distances from the edge of the gel interface of all cells invading into the collagen gel were measured. **G,** The same assay as in (**F**) was performed in the presence of different concentrations of collagen I. **H,** S2-CP8 cells stably expressing ARL4C-tdTomato were observed with time-lapse imaging. Arrowheads indicate the tips of cell protrusions. The region in the yellow dashed squares is shown enlarged in the bottom image. Fluorescence intensities of the cytoplasm and cell protrusions were measured and plotted as a function of time. **I,** S2-CP8 cells expressing ARL4C-tdTomato were subjected to a 3D collagen I gel invasion assay with ^DQ^collagen I, and stained with phalloidin and Hoechst 33342. The regions in the yellow dashed squares (A, protrusion; B, cell body) are enlarged. **J,** S2-CP8 cells transfected with control ASO or ARL4C ASO-1316 were subjected to a 3D collagen I gel invasion assay with ^DQ^collagen I. The percentages of cells with ^DQ^collagen I-positive protrusions compared with the total number of cells were calculated. **A–D,F,G,J,** Data are shown as the mean ± s.d. of 3 independent experiments. *P* values were calculated using a two-tailed Student’s t-test. Scale bars in **F,G,** 100 μm; **H,** 20 μm; **I,** 10 μm; **J,** 5 μm. n.s. not significant. *, *P* < 0.05; **, *P* < 0.01. See Figure 2-source data 1.

For visualization of cancer cells invading through the extracellular matrix (ECM)(Poincloux et al, 2009), a 3D microfluidic cell culture with type I collagen(Farahat et al, 2012; Shin et al, 2012) (3D gel invasion assay) was performed (Figure 2E). S2-CP8 cells invaded into type I collagen, and individual cells formed protrusions at the leading side of the cells (Figure 2F). In contrast, ARL4C knockdown decreased invasive ability (Figure 2F). When the collagen concentration was reduced, S2-CP8 cells invaded irrespective of ARL4C knockdown (Figure 2G), suggesting that their invasive ability is not required for cells to move into the ECM when collagen fiber-formed 3D net structures are sparse. Furthermore, in the 3D gel invasion assay, membrane protrusions were time-dependently observed in the direction of invasion, and ARL4C-tdTomato accumulated in the tips of membrane protrusions (Figure 2H; Figure 2-video 1).

To visualize the relationship between the localization of ARL4C and matrix degradation, the steady-state activity of cell-derived collagenase was measured as the dequenched signal emitted from collagen I fibers with dye-quenched (DQ) FITC (^DQ^collagen I)(Wolf et al, 2007) in the 3D gel invasion assay. Collagenase-induced fluorescence dequenching was detected in the collagen fibers crossing the tips of the protrusions but not in the cell body (Figure 2I). Collagenase activity was decreased when ARL4C was depleted (Figure 2J), suggesting that ARL4C is involved in degradation of the ECM through its localization to the tips of cell protrusions.

While cell protrusions are suggested to be involved in invasion, invadopodia are well-known membrane protrusions that localize at the ventral surfaces of cells and are active in ECM degradation during cancer invasion(Murphy & Courtneidge, 2011). To analyze invadopodia, the cells were grown on gelatin-coated glass coverslips (Figure 2-figure supplement 2A). Dark areas represent gelatinolytic activity of invadopodia and are equal to invadopodia structures. BxPC-3 cells, which expressed low levels of ARL4C, exhibited invadopodia clearly, whereas S2-CP8 and PANC-1 cells, which highly express ARL4C, did not (Figure 2-figure supplement 2A). It is notable that S2-CP8 and PANC-1 cells formed membrane protrusions but BxPC-3 did not. Overexpression of ARL4C-GFP in BxPC-3 cells (BxPC-3/ARL4C-GFP cells) did not affect the numbers of invadopodia but did promote invasive ability (Figure 2-figure supplement 2B-D). Wild-type BxPC-3 cells formed a round shape in 3D culture conditions, whereas BxPC-3/ARL4C-GFP cells formed membrane protrusions and ARL4C-GFP was observed at the tips of the membrane protrusions (Figure 2-figure supplement 2E and F), suggesting that ARL4C expression is not required for invadopodia formation. Taken together, ARL4C is localized to membrane protrusions and plays an important role in the invasion of pancreatic cancer cells.

### IQGAP1 is an ARL4C-interacting protein

ARL4C recruits cytohesin2 to the plasma membrane through their direct interaction in HeLa cells(Hofmann et al, 2007). In S2-CP8 cells, ARL4C did not bind to cytohesin2 (Figure 3-figure supplement 1A), and knockdown of cytohesin2 had no effect on the migratory or invasive ability (Figure 3-figure supplement 1B). Furthermore, cytohesin2 was distributed throughout the cytosol in S2-CP8 cells, whereas it was localized to the cell periphery of HeLaS3 cells (Figure 3-figure supplement 1C). Whereas ARL4C ASO inhibited RAC1 activity in A549 cells(Fujii et al, 2015), the ASO did not affect RAC1 activity in S2-CP8 cells and overexpression of ARL4C did not affect it in BxPC-3 cells (Figure 3-figure supplement 1D). Although ARL4C induces the nuclear import of YAP/TAZ in HCT116 cells(Harada et al, 2019), ARL4C knockdown did not inhibit it in pancreatic cancer cells (Figure 3-figure supplement 1E). These results suggest that cytohesin2 neither functions downstream of ARL4C nor is involved in migration or invasion of S2-CP8 cells and prompted us to explore an uncharacterized effector protein of ARL4C.

ARL4C-FLAG-HA–binding proteins were precipitated and the precipitates were analyzed by mass spectrometry (Figure 3A). Among the possible interacting proteins, IQGAP1 was further studied (Figure 3A; Supplementary file 1 Table 2) because its expression is associated with the aggressiveness of various types of cancer(Johnson et al, 2009). Ectopically expressed and endogenous ARL4C were associated with endogenous IQGAP1 in S2-CP8 cells (Figure 3B and C). ARL4C-FLAG-HA and ARL4C^Q72L^-FLAG-HA formed a complex with GFP-IQGAP1 to the similar levels, but ARL4C^T27N^-FLAG-HA showed diminished binding to GFP-IQGAP1 in X293T cells (Figure 3D).

**Figure 3.**
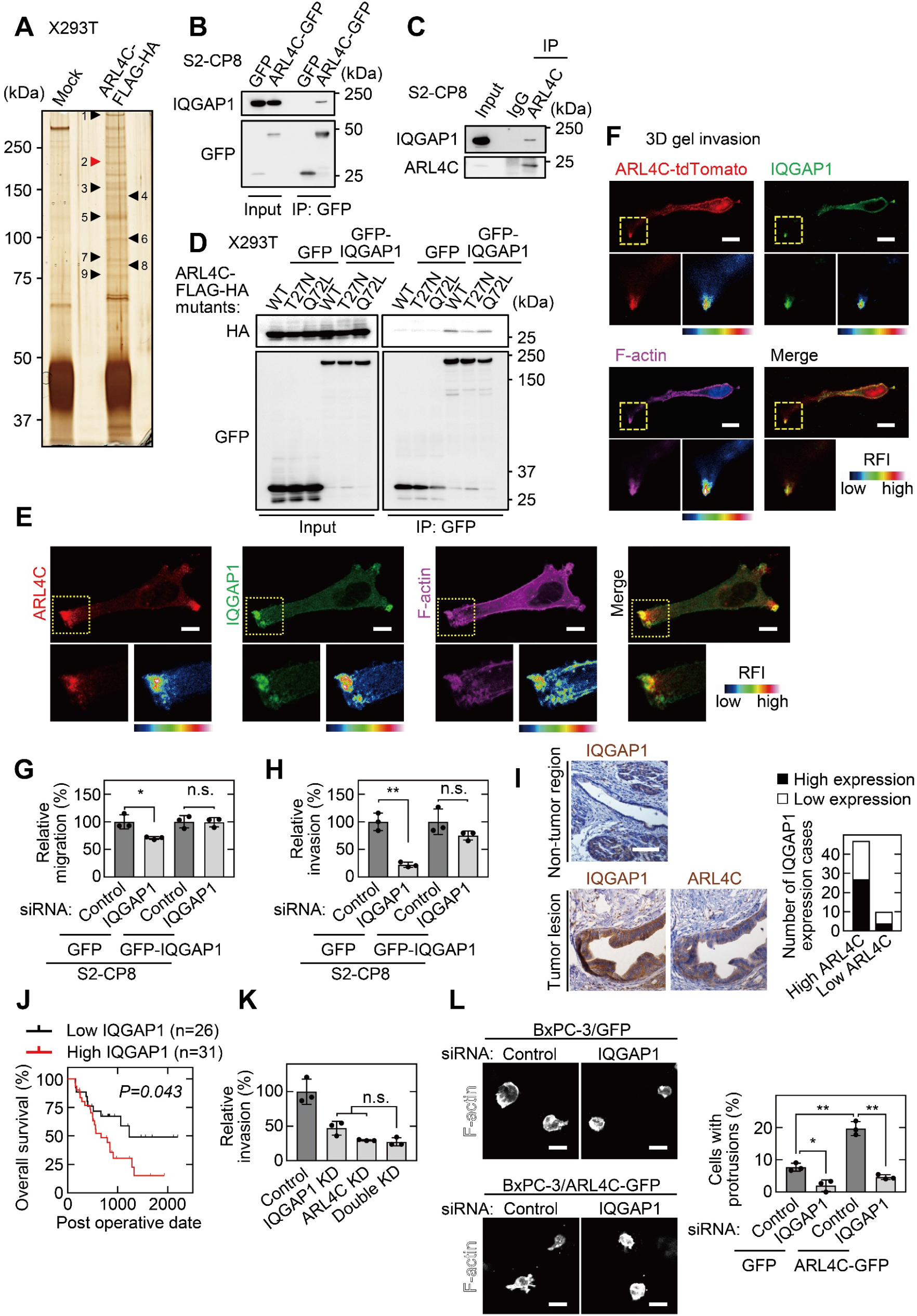
IQGAP1 is a novel ARL4C-interacting protein. **A,** The ARL4C-interacting proteins in X293T cells were analyzed by mass spectrometry. The results are listed in Table EV2. Arrowheads indicate the identified proteins, including IQGAP1 (red). **B,C,** Lysates of S2-CP8 cells expressing ARL4C-GFP (**B**) or S2-CP8 WT cells (**C**) were immunoprecipitated with anti-GFP antibody (**B**) or anti-ARL4C antibody (**C**), and the immunoprecipitates were probed with the indicated antibodies. **D,** Lysates of X293T cells expressing the indicated proteins were immunoprecipitated with anti-GFP antibody, and the immunoprecipitates were probed with the indicated antibodies. **E,** S2-CP8 cells were stained with the indicated antibodies. Images of ARL4C and IQGAP1 were merged. The regions in the yellow dashed squares are shown enlarged in the left bottom images. The right bottom images are shown with a false color representation of fluorescence intensity. More than 50 cells were imaged and the representative image is shown. **F,** S2-CP8 cells expressing ARL4C-tdTomato were subjected to a 3D collagen I gel invasion assay and were stained with the indicated antibodies. Images of ARL4C and IQGAP1 were merged. Enlarged images of the regions in the yellow dashed squares are shown in a false color representation of fluorescence intensity on the right. **G,H,** S2-CP8 cells expressing GFP or GFP-IQGAP1 were transfected with the indicated siRNAs and subjected to migration (**G**) and invasion (**H**) assays. Migratory and invasive abilities are expressed as the percentage of the same cells transfected with control siRNA. **I,** PDAC tissues were stained with the indicated antibodies and hematoxylin. IQGAP1 expression cases in high or low ARL4C expression lesions are shown. **J,** The relationship between overall survival and IQGAP1 expression in PDAC patients was analyzed. **K,** S2-CP8 cells depleted of the indicated proteins were subjected to an invasion assay. Invasive activities are expressed as the percentage of control cells. **L,** BxPC-3 cells stably expressing GFP or ARL4C-GFP were transfected with the indicated siRNAs and then cultured in 3D collagen I gel. The percentages of cells with protrusions compared with the total number of cells were calculated. **G,H,K,L,** Data are shown as the mean ± s.d. of 3 independent experiments. *P* values were calculated using a two-tailed Student’s t-test (**G,H**) or one-way ANOVA followed by Bonferroni post hoc test (**K,L**). **J,** The data were analyzed by Kaplan-Meier survival curves, and a log-rank test was used for statistical analysis. **E,F,** False color representations were color-coded on the spectrum. Scale bars in **E,** 10 μm; **F,** 20 μm; **I,L,** 50 μm. KD, knockdown. RFI, relative fluorescence intensity. n.s., not significant. *, *P* < 0.05; **, *P* < 0.01. See Figure 3-source data 1.

Using another anti-ARL4C antibody for the immunocytochemical study (Figure 3-figure supplement 2A and B), ARL4C and IQGAP1 were shown to accumulate to membrane protrusions at endogenous level in S2-CP8 and PANC-1 cells under Matrigel-coated 2D culture conditions (Figure 3E; Figure 3-figure supplement 2C). Colocalization of ARL4C and IQGAP1 at membrane protrusions was observed in 94% of cells with ARL4C accumulation to the protrusions. In 3D culture conditions, IQGAP1 was found at the tips of membrane protrusions, similar to ARL4C-tdTomato (Figure 3F). IQGAP1 siRNA inhibited the migratory and invasive abilities in S2-CP8 and PANC-1 cells, and the cells expressing GFP-IQGAP1 expression were resistant to IQGAP1 siRNA (Figure 3G and H; Figure 3-figure supplement 2D).

IQGAP1 was highly expressed in 31 of 57 PDAC patients (54%), whereas it was minimally detected in non-tumor regions of pancreatic ducts (Figure 3I). The anti-IQGAP1 antibody was validated by Western blotting and immunocytochemical and immunohistochemical analyses (Figure 3-figure supplement 2E-G). Although higher expression of IQGAP1 was not associated with clinical parameters (Supplementary file 1 Table 3), IQGAP1 expression correlated with decreased overall survival (Figure 3J). Similar results were obtained from the analysis of TCGA and GTEx datasets (Figure 3-figure supplement 2H and I). Of 47 PDAC patients with high ARL4C expression, IQGAP1 was highly expressed in 27 patients (Figure 3I). Higher expression of ARL4C in the patients positive for IQGAP1 was associated with perineural invasion (Supplementary file 1 Table 4). The overall survival of PDAC patients who were double positive for ARL4C and IQGAP1 tended to be worse (Figure 3-figure supplement 2J).

Simultaneous knockdown of ARL4C and IQGAP1 decreased the invasive ability, but the inhibitory degree was similar to that induced by knockdown of either ARL4C or IQGAP1 (Figure 3K). IQGAP1 knockdown inhibited ARL4C-induced formation of membrane protrusions in BxPC-3 cells cultured under 3D conditions (Figure 3L). Thus, IQGAP1 functions downstream of ARL4C and they regulate invasion in identical signaling pathways.

### The polybasic region of ARL4C is required for its binding to IQGAP1

ARL4C is modified by myristate at the N terminus and has a polybasic region (PBR), comprising nine Lys or Arg residues, at the C terminus(Donaldson & Jackson, 2011). ARL4C^G2A^, whose N-terminal myristoylation site (Gly2) is mutated to Ala, and ARL4C^ΔPBR^ were expressed in S2-CP8 cells. In contrast to ARL4C-GFP, ARL4C^G2A^-GFP and ARL4C^ΔPBR^-GFP were not accumulated at membrane protrusions but distributed throughout the cytosol (Figure 4A and B), and both mutants severely decreased the binding activity to GFP-IQGAP1 (Figure 4C). The C-terminal region of KRAS includes the PBR and the CAAX motif, which is farnesylated, and fusion of the KRAS C-terminal region triggers the localization of the proteins to the cell surface membrane(Hancock et al, 1990). The KRAS C-terminal region was fused to the ARL4C mutants, which were referred to as ARL4C-GFP-Cterm. Both ARL4C^G2A^-GFP-Cterm and ARL4C^ΔPBR^-GFP-Cterm were localized to membrane protrusions (Figure 4B). However, although ARL4C^G2A^-FLAG-HA-Cterm formed a complex with GFP-IQGAP1, ARL4C^ΔPBR^-FLAG-HA-Cterm did not (Figure 4D), suggesting that membrane localization of ARL4C is not sufficient for its binding to IQGAP1.

**Figure 4.**
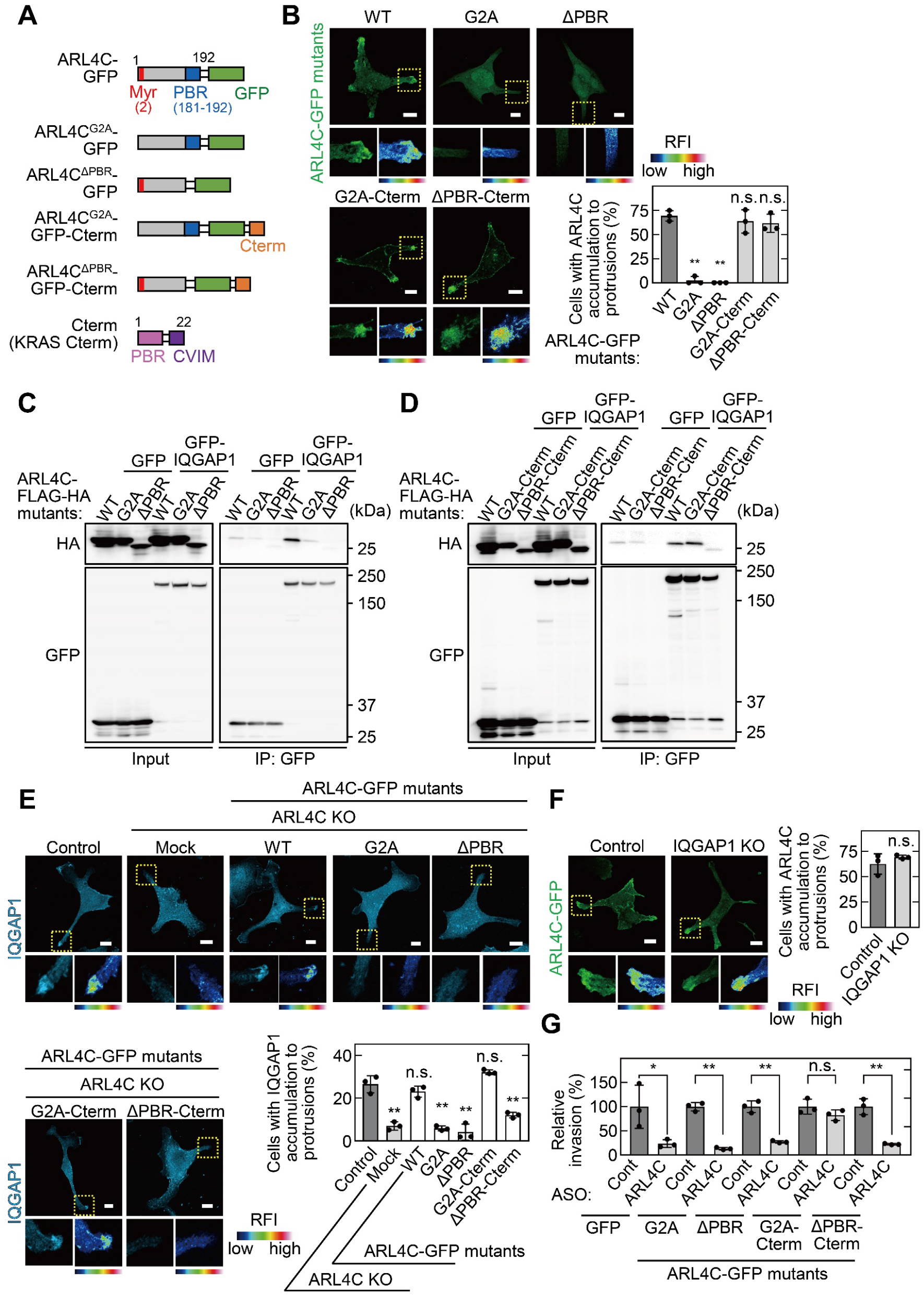
The PBR of ARL4C is required for ARL4C and IQGAP1 binding. **A,** A schematic representation of four ARL4C-GFP mutants is shown. **B,** S2-CP8 cells were transfected with the indicated mutants of ARL4C-GFP. The percentages of cells with ARL4C-GFP mutant accumulated at membrane protrusions compared with the total number of cells were calculated. **C,D,** Lysates of X293T cells expressing the indicated proteins were immunoprecipitated with anti-GFP antibody and the immunoprecipitates were probed with anti-HA and anti-GFP antibodies. **E,** S2-CP8 WT or ARL4C KO cells transfected with control or the indicated mutants of ARL4C-GFP were stained with anti-IQGAP1 antibody. The percentages of cells with IQGAP1 accumulated at membrane protrusions compared with the total number of cells were calculated. **F,** S2-CP8 WT or IQGAP1 KO cells were transfected with ARL4C-GFP. The percentages of cells with ARL4C-GFP accumulated at membrane protrusions compared with the total number of cells were calculated. **G,** S2-CP8 cells stably expressing GFP or the indicated mutants of ARL4C-GFP were transfected with control or ARL4C ASO and subjected to invasion assays. Invasive ability is expressed as the percentage of the same cells transfected with control ASO. **B,E–G,** Data are shown as the mean ± s.d. of 3 independent experiments. *P* values were calculated using a two-tailed Student’s t-test (**F,G**) or one-way ANOVA followed by Bonferroni post hoc test (**B,E**). **B,E,F,** The regions in the yellow dashed squares are shown enlarged in the left bottom images. The right bottom images are shown in a false color representation of fluorescence intensity. False color representations were color-coded on the spectrum. Scale bars in **B,E,F,** 10 μm. KO, knockout. RFI, relative fluorescence intensity. n.s., not significant. *, *P* < 0.05; **, *P* < 0.01. See Figure 4-source data 1.

The localization of IQGAP1 to membrane protrusions was lost in ARL4C knock out (KO) cells but not vice versa (Figure 4E and F). In ARL4C KO cells, ARL4C-GFP and ARL4C^G2A^-GFP-Cterm rescued the recruitment of IQGAP1 to the plasma membrane, unlike ARL4C^G2A^-GFP, ARL4C^ΔPBR^-GFP, and ARL4C^ΔPBR^-GFP-Cterm (Figure 4E). Therefore, for IQGAP1 to be recruited to membrane protrusions, the localization of ARL4C to the plasma membrane through the PBR might be necessary. In addition, ARL4C ASO-1316 inhibition of invasive ability was cancelled by expression of ARL4C^G2A^-GFP-Cterm but not by that of ARL4C^G2A^-GFP, ARL4C^ΔPBR^-GFP, or ARL4C^ΔPBR^-GFP-Cterm (Figure 4G; Figure 4-figure supplement 1A). Thus, the binding of ARL4C and IQGAP1 in membrane protrusions could be essential for the invasive ability.

### ARL4C recruits IQGAP1 to membrane protrusions in a PI(3,4,5)P3-dependent manner

PI(4,5)P2 (PIP2) and PI(3,4,5)P3 (PIP3) are required for ARL4C membrane targeting(Heo et al, 2006). The pleckstrin homology (PH) domain functions as a protein- and phospholipid-binding structural protein module(Maffucci & Falasca, 2001). The PH domains of PLCδ and GRP1 prefer to bind to PIP2 and PIP3, respectively(Lemmon, 2008). GFP-PLCδ^PH^ was detected throughout the cell surface membrane, whereas GFP-GRP1^PH^ was accumulated in membrane protrusions (Figure 5A).

**Figure 5.**
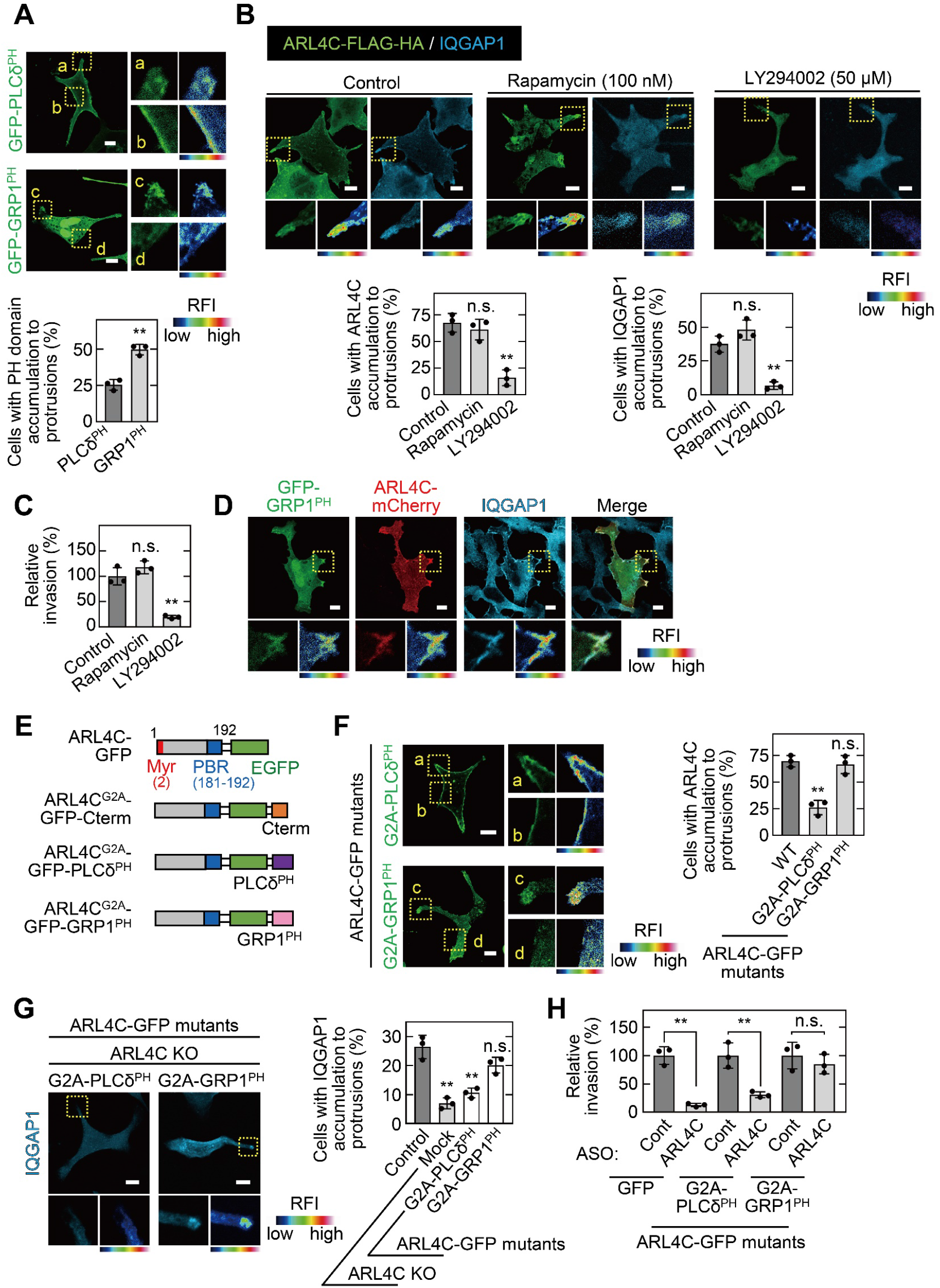
ARL4C recruits IQGAP1 to membrane protrusions in a PIP3-dependent manner. **A,** S2-CP8 cells were transfected with GFP-PLCδ^PH^ or GFP-GRP1^PH^. The percentages of cells with GFP-PLCδ^PH^ or GFP-GRP1^PH^ accumulated at membrane protrusions compared with the total number of cells were calculated. **B,** S2-CP8 cells expressing FRB-CFP, mRFP-FKBP-5-ptase domain, and ARL4C-FLAG-HA were treated with or without rapamycin or LY294002 and stained with anti-HA and anti-IQGAP1 antibodies. The percentages of cells with IQGAP1 or ARL4C-FLAG-HA accumulated at membrane protrusions compared with the total number of cells were calculated. **C**, S2-CP8 cells expressing FRB-CFP and mRFP-FKBP-5-ptase domain were treated with or without rapamycin or LY294002 and subjected to an invasion assay. Invasive activities are expressed as the percentage of control cells. **D,** S2-CP8 cells expressing ARL4C-mCherry and GFP-GRP1^PH^ were stained with anti-IQGAP1 antibody. Images of GFP-GRP1^PH^, ARL4C-mCherry, and IQGAP1 were merged. **E,** A schematic representation of ARL4C-GFP mutants is shown. **F,** S2-CP8 cells were transfected with the indicated mutants of ARL4C-GFP. The percentages of cells with ARL4C-GFP mutant accumulated at membrane protrusions compared with the total number of cells were calculated. **G,** ARL4C KO cells expressing control or the indicated mutants of ARL4C-GFP were stained with anti-IQGAP1 antibody. Quantification was performed as in (**B**). **H,** S2-CP8 cells stably expressing GFP or the indicated mutants of ARL4C-GFP were transfected with control or ARL4C ASO and subjected to an invasion assay. Invasive ability is expressed as the percentage of the same cells transfected with control ASO. **A,F,** Enlarged images of the regions in the yellow dashed squares and a false color representation of fluorescence intensity are shown on the right. (a) and (c) show the protrusion, and (b) and (d) show the cell body. **B,D,G,** The regions in the yellow dashed squares are shown enlarged in the left bottom images. The right bottom images are shown in a false color representation of fluorescence intensity. **A–C,F–H,** Data are shown as the mean ± s.d. of 3 independent experiments. *P* values were calculated using a two-tailed Student’s t-test (**A,H**) or one-way ANOVA followed by Bonferroni post hoc test (**B,C,F,G**). **A,B,D,F,G,** False color representations were color-coded on the spectrum. Scale bars in **A,B,D,F,G**, 10 μm. KO, knockout. RFI, relative fluorescence intensity. n.s., not significant. **, *P* < 0.01. See Figure 5-source data 1.

The levels of PIP2 and PIP3 in the plasma membrane were decreased by a rapamycin-inducible PIP2-specific phosphatase (Inp54p)(Suh et al, 2006) and a PI3 kinase inhibitor LY294002(Petrie et al, 2012), respectively. PIP3 depletion decreased the membrane targeting of ARL4C and IQGAP1 and reduced the invasive ability, but PIP2 depletion did not (Figure 5B and C). IQGAP1 and ARL4C-mCherry colocalized with GRP1^PH^ in membrane protrusions (Figure 5D), suggesting that both proteins accumulate in the cell peripheral regions containing PIP3 and promote invasion.

To reveal the importance of PIP3 for the localization area of ARL4C and IQGAP1, PLCδ^PH^ or GRP1^PH^ was fused to the C terminus of ARL4C^G2A^-GFP (Figure 5E). While both ARL4C^G2A^-GFP-GRP1^PH^ and ARL4C^G2A^-GFP-PLCδ^PH^ formed a complex with GFP-IQGAP1, the former construct was localized to membrane protrusions, but the latter construct was present throughout the cell surface membrane (Figure 5F; Figure 5-figure supplement 1A). Consistently, in ARL4C KO cells, the localization of IQGAP1 to membrane protrusions was rescued by ARL4C^G2A^-GFP-GRP1^PH^ but not by ARL4C^G2A^-GFP-PLCδ^PH^ (Figure 5G). Furthermore, ARL4C ASO-1316 inhibited the invasive ability of S2-CP8 cells expressing ARL4C^G2A^-GFP-PLCδ^PH^ but not those expressing ARL4C^G2A^-GFP-GRP1^PH^ (Figure 5H; Figure 5-figure supplement 1B). Taken together, these results suggest that PIP3-dependent membrane targeting of ARL4C recruits IQGAP1 to membrane protrusions and promotes invasion.

### ARL4C is involved in the focal delivery of MMP14 to membrane protrusions through IQGAP1

IQGAP1 is involved in the trafficking of MMP14-containing vesicles to invasive protrusions of cancer cells(Sakurai-Yageta et al, 2008). TCGA dataset showed that expression of *MMP14* mRNA in pancreatic cancer patients is positively correlated with that of both *ARL4C* and *IQGAP1* mRNA (Figure 6-figure supplement 1A). In addition, MMP14 expression was associated with poor prognosis (Figure 6-figure supplement 1B).

Cell surface MMP14-GFP accumulated in membrane protrusions containing IQGAP1 and ARL4C-FLAG-HA (Figure 6A). MMP14-GFP disappeared from the membrane protrusions of ARL4C KO and IQGAP KO cells and the phenotype was rescued by expression of ARL4C-FLAG-HA and FLAG-HA-IQGAP1 (Figure 6B and C). The failure of MMP14 membrane targeting in ARL4C KO cells was rescued by expression of ARL4C^G2A^-FLAG-HA-Cterm but not by that of ARL4C^G2A^-FLAG-HA, ARL4C^ΔPBR^-FLAG-HA, or ARL4C^ΔPBR^-FLAG-HA-Cterm (Figure 6B). In addition, PIP3 depletion, but not PIP2 depletion, suppressed the membrane localization of MMP14 (Figure 6D). Therefore, in co-operation with ARL4C and IQGAP1, MMP14 is likely to be trafficked to membrane protrusions with PIP3 accumulation.

**Figure 6.**
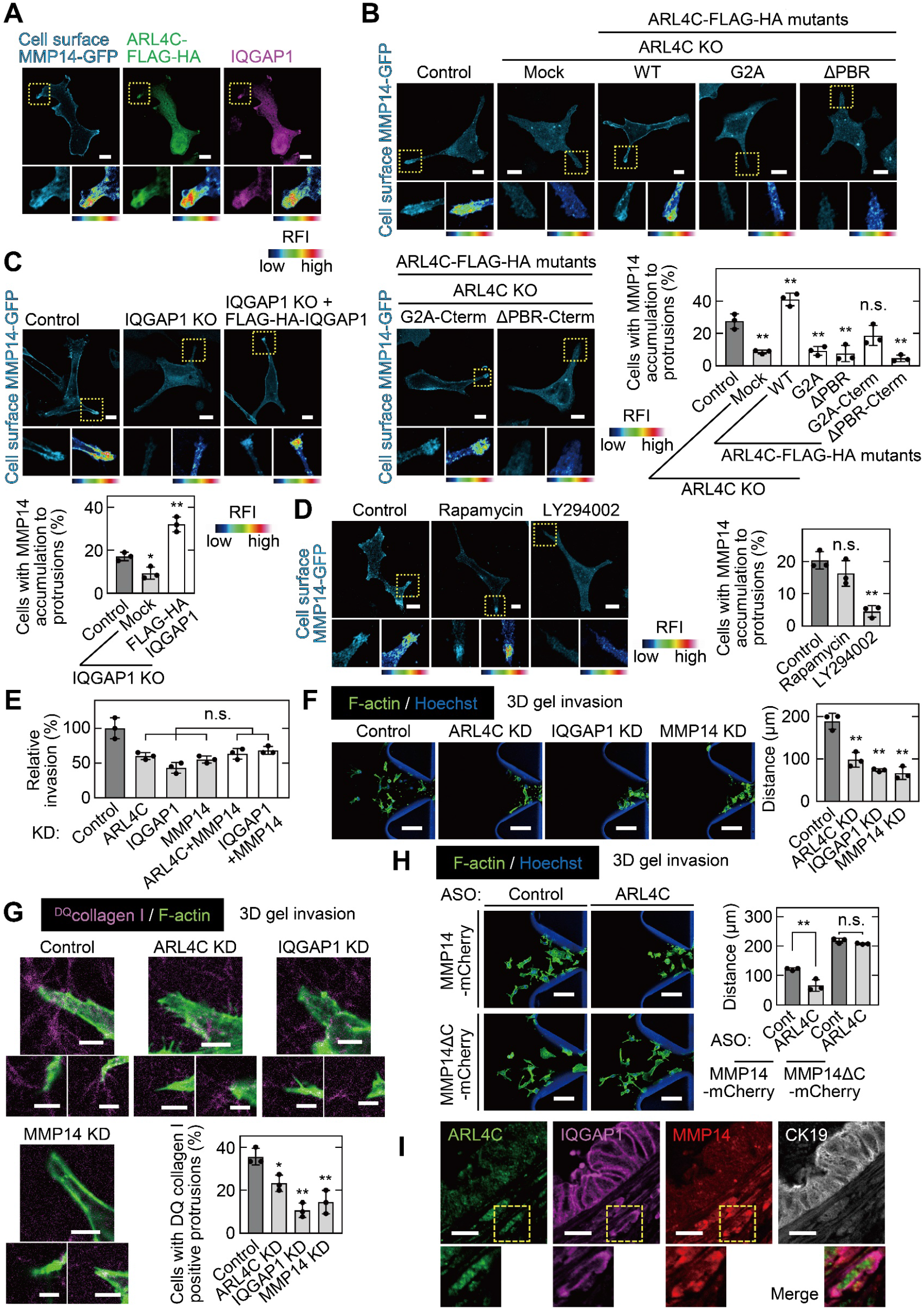
ARL4C is involved in focal delivery of MMP14 to membrane protrusions through IQGAP1. **A,** S2-CP8 cells expressing MMP14-GFP and ARL4C-FLAG-HA were stained with anti-MMP14 without permeabilization, followed by permeabilization and staining with anti-HA and anti-IQGAP1 antibodies. **B,** S2-CP8 WT or ARL4C KO cells expressing MMP14-GFP and the indicated mutants of ARL4C-FLAG-HA were stained with anti-MMP14 without permeabilization. The percentages of cells with MMP14 accumulated at membrane protrusions compared with the total number of cells were calculated. **C,** The same assay as in (**B**) was performed except with S2-CP8 WT or IQGAP1 KO cells expressing MMP14-GFP and FLAG-HA-IQGAP1. **D,** S2-CP8 cells expressing MMP14-GFP, FRB-CFP, and mRFP-FKBP-5-ptase domain were treated with 100 nM rapamycin or 50 μM LY294002 for 30 min. Staining and quantification were performed as in (**B**). **E,** S2-CP8 cells depleted of the indicated proteins were subjected to an invasion assay. Invasive activities are expressed as the percentage of control cells. **F-H,** S2-CP8 cells (**F,G**) or S2-CP8 cells expressing MMP14-mCherry or MMP 14ΔC-mCherry (**H**) depleted of the indicated proteins were subjected to a 3D collagen I gel invasion assay with ^DQ^collagen I. The distances from the edge of the gel interface of all cells that invaded into the gel were measured (**F,H**). The percentages of cells with ^DQ^collagen I-positive protrusions compared with the total number of cells were calculated. (**G**). **I,** PDAC tissues were stained with the indicated antibodies. Images of ARL4C, IQGAP1, and MMP14 were merged. Magnified fluorescence images are shown in the bottom panels. Nine patient samples were imaged and the representative images are shown. **A–D,** The regions in the yellow dashed squares are shown enlarged in left bottom and a false color representation of fluorescence intensity is shown in right bottom. False color representations were color-coded on the spectrum. **B–H,** Data are shown as the mean ± s.d. of 3 independent experiments. *P* values were calculated using a two-tailed Student’s t-test (**H**) or one-way ANOVA followed by Bonferroni post hoc test (**B–G**). Scale bars in **A–D,** 10 μm; **F,H,** 100 μm; **G,** 5 μm; **I,** 20 μm. KO, knockout; KD, knockdown. RFI, relative fluorescence intensity. n.s., not significant. *, *P* < 0.05; **, *P* < 0.01. See Figure 6-source data 1.

Consistent with these results, the inhibited invasive ability after double knockdown of ARL4C and MMP14 or IQGAP1 and MMP14 by siRNA was similar to that seen after single knockdown of ARL4C, IQGAP1, or MMP14 (Figure 6E; Figure 6-figure supplement 1C and D). Knockdown of ARL4C, IQGAP1, or MMP14 also decreased invasive ability in 3D microfluidic cell culture (Figure 6F) and the collagenase activity was also reduced (Figure 6G). Previous work has shown that MMP14^ΔC^(Δ563–582) lacking the cytoplasmic region fails to be endocytosed(Jiang et al, 2001). Here, MMP14^ΔC^ was retained in membrane protrusions of ARL4C-depleted cells (Figure 6-figure supplement 1E), and the ARL4C knockdown-mediated decreases in cell invasion and collagen degradation were rescued by MMP14^ΔC^ (Figure 6H; Figure 6-figure supplement 1F and G). Thus, ARL4C-dependent recruitment of MMP14 to membrane protrusions is required for cell invasion.

MMP14 was detected in similar lesions to ARL4C and IQGAP1 in serial PDAC specimens (Figure 6-figure supplement 1H). Notably, a group of cells invaded the surrounding interstitial tissues, and concurrently expressed ARL4C, IQGAP1, and MMP14 (Figure 6I; Figure 6-figure supplement 1H). Taken together, these results support the idea that the ARL4C–IQGAP1–MMP14 signaling axis participates in pancreatic cancer cell invasion.

### ARL4C ASO inhibits pancreatic tumor metastasis *in vivo*

To show that ARL4C is indeed involved in cancer cell invasion *in vivo*, the effects of subcutaneous injection of ARL4C ASO-1316 on an orthotopic transplantation model was tested. S2-CP8 cells expressing luciferase were injected into the pancreas of nude mice, and control ASO or ARL4C ASO-1316 was subcutaneously injected from day 3 (Figure 7A). After 2 and 3 weeks, ARL4C ASO-1316 suppressed the luminescence signal compared with control ASO (Figure 7B). Whereas ARL4C ASO-1316 did not reduce the size of the primary tumor in the pancreas, the ASO decreased the numbers of lymph node metastases and tended to improve the survival (Figure 7C and D; Figure 7-figure supplement 1A).

**Figure 7.**
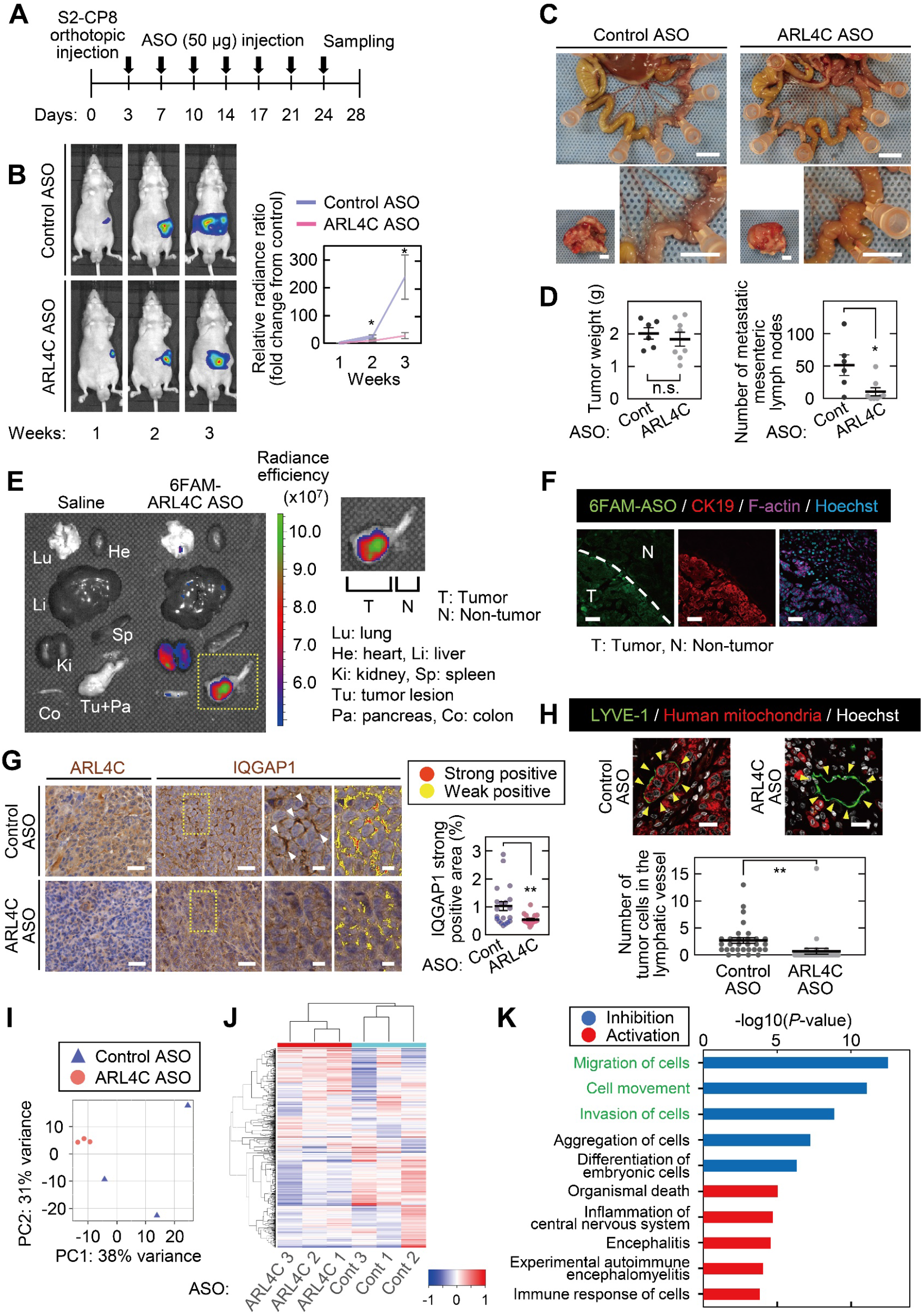
ARL4C ASO inhibits pancreatic tumor invasion *in vivo*. **A,** S2-CP8/Luciferase cells were implanted into the pancreas of nude mice, and control ASO (n = 6) or ARL4C ASO-1316 (n = 7) was subcutaneously administered. **B,** Bioluminescence images of the intraperitoneal tumors are presented (left) and quantification of the tumor burden is shown (right). The data are presented as the mean ± s.e.m. of the fold change in luminescent intensity relative to that of week 1 treated with control ASO. **C,** Representative images of the tumors in the pancreas (left bottom) and metastatic mesenteric lymph nodes (top and right bottom) are shown. **D,** Primary tumor weight (left) and metastatic mesenteric lymph node number are presented (right). Data are shown as the mean ± s.e.m. **E,F,** 4 h after subcutaneous injection of 6-FAM-ARL4C ASO-1316 into tumor-bearing mice, the fluorescence intensities of various organs were measured (**E**), and the sections prepared from the pancreas were stained with the indicated antibodies (**F**). Area indicated by yellow dashed square is enlarged on the right panel (**E**). **G,** Sections from the pancreatic tumor were stained with the indicated antibodies and hematoxylin. The two panels on the right show enlarged images of the yellow dashed squares. Positive staining of IQGAP1 is color-coded as yellow (weakly positive) or red (strongly positive). The percentage of the strongly positive IQGAP1 area was calculated. Data are shown as the mean ± s.e.m. Twenty fields were analyzed from 3 mice per group. **H,** Sections from the pancreatic tumor were stained with the indicated antibodies. The numbers of tumor cells in the lymphatic vessels (indicated with arrowheads) were counted. Data are shown as the mean ± s.e.m. Thirty lymphatic vessels were analyzed from 3 mice per group. **I,J,** RNA sequencing was performed for S2-CP8-derived primary tumors, and the results of principal component analysis (**I**) and hierarchical clustering (**J**) are shown. **K,** Differentially expressed genes were subjected to Ingenuity Pathway Analysis (IPA). The top five disease or function annotations of the positive and negative Z-score groups are shown. Bars indicate the -log(*P* value). Inhibited pathways are represented by blue-colored bars while activated pathways are shown by red-colored bars. **B, D,G,H,** *P* values were calculated using a two-tailed Student’s t-test. Scale bars in **C,** 5 mm; **F,G,** 50 μm; **H,** 20 μm. n.s., not significant. *, *P* < 0.05; **, *P* < 0.01. See Figure 7-source data 1.

When 6-FAM–labeled ARL4C ASO-1316 was subcutaneously injected into tumor-bearing mice, the fluorescence was specifically detected in the pancreas (Figure 7E). 6-FAM–labeled ARL4C ASO-1316 was highly accumulated in tumor lesions but not in the neighboring normal tissues (Figure 7F), indicating that ASO was incorporated into tumor lesions after systemic injection. In primary pancreatic tumors, ARL4C ASO-1316 reduced ARL4C expression and decreased the localization of IQGAP1 to the cell surface area (Figure 7G; Figure 7-figure supplement 1B). Tumor cells were observed in lymphatic vessels of peritumoral areas of control ASO-treated mice but not in those of ARL4C ASO-treated mice (Figure 7H).

To compare molecular characteristics between pancreatic tumors from mice injected with control ASO and ARL4C ASO-1316, RNA sequence analysis was performed for primary tumors. Principal component analysis (PCA) indicated a clear difference in the gene expression profiles of tumors from control ASO- and ARL4C ASO-1316–treated mice (Figure 7I). Furthermore, hierarchical clustering revealed a drastic change in expression of genes due to ARL4C ASO-1316 injection (Figure 7J). Two hundred and three differentially expressed genes (DEGs) were detected, and by subjecting them to Ingenuity Pathway Analysis (IPA), the top 5 significantly enriched terms of the biological process of molecular function in the inhibition and activation of the pathways were obtained (Figure 7K). In particular, DEGs linked to the inhibition of the pathways in ARL4C ASO-1316–treated mice were predicted to be involved in terms such as cell migration and invasion (Figure 7K). Taken together, these results suggest that ARL4C ASO inhibits the invasion of tumor cells into lymphatic vessels *in vivo,* and the gene profiles of tumors treated with ARL4C ASO *in vivo* support the putative functions of ARL4C in pancreatic cancer invasion.

## Discussion

Pancreatic cancer represents one of the leading causes of cancer death, despite advances in cancer therapy(Keleg et al, 2003). Major problem of pancreatic cancer is uncontrollable invasion and metastasis. In this study, we found that the ARL4C–IQGAP1–MMP14 signaling axis is involved in pancreatic cancer invasion. Because ARL4C expression is induced by Wnt and EGF signaling, it is reasonable that ARL4C would be expressed in a β-catenin– and RAS-dependent manner in pancreatic cancer cells. ARL4C is a unique small G protein because it is constitutively active, regardless of wild-type(Burd et al, 2004; Matsumoto et al, 2017). The long interswitch region of ARL4C may prevent the retractile conformation change in the GDP-bound state(Burd et al, 2004; Pasqualato et al, 2002). ARL4C could be a constitutively active form without active mutations, and its activity may be controlled by transcriptional regulation.

ARL4C binds to cytohesin2(Hofmann et al, 2007), leading to activation of ARF6–RAC–RHO– YAP/TAZ signaling in colon and lung cancer cells(Fujii et al, 2015; Kimura et al, 2020). Because ARL4C did not bind to cytohesin2 but to IQGAP1 in pancreatic cancer cells, it is likely that ARL4C regulates different downstream signaling pathways in a cancer cell context-dependent manner. Invadopodia are the unique structures observed at the ventral sites of certain types of cancer cells(Dalaka et al, 2020; Murphy & Courtneidge, 2011). However, in S2-CP8 and PANC-1 cells highly expressing ARL4C, invadopodia were not formed and ARL4C was observed in membrane protrusions. Because both structures are formed by similar molecules, including IQGAP1 and MMP14(Caswell & Zech, 2018; Jacquemet et al, 2013), ARL4C may determine the delivery of signaling components and cellular machineries to the cell peripheral membrane.

Both myristoylation and the PBR of ARL4C support plasma membrane targeting(Heo et al, 2006). In our results, both motifs were necessary for the localization of ARL4C to the plasma membrane, whereas the PBR, rather than myristoylation, was indispensable for the activity of the ARL4C–IQGAP1–MMP14 signaling axis. Phosphoinositides have been implicated in many aspects of cell physiology(Di Paolo & De Camilli, 2006). PIP3 is localized to the leading edge of migrating cells and invadopodia of cancer cells(Saykali & El-Sibai, 2014) and recruits cytosolic proteins containing lipid-binding domains, such as the PH domain, to the plasma membrane(Toker & Cantley, 1997). ARL4C in pancreatic cancer cells preferred PIP3 to PIP2. Because PI3 kinase is one of the direct effector proteins of RAS(Castellano & Downward, 2011; Rodriguez-Viciana et al, 1994), RAS-dependent PI3 kinase activation and ARL4C expression could co-operatively function to promote pancreatic cancer invasion.

In conclusion, this study clarified that invasion of pancreatic cancer cells is promoted by ARL4C, which is induced by KRAS and Wnt signaling, and association of ARL4C with IQGAP1 and MMP14 at the membrane protrusion is essential for the invasive ability. The novel functions of ARL4C were confirmed by the mouse model. The inhibition of ARL4C expression by ARL4C ASO could directly inhibit invasion ability of pancreatic cancer cells and may indirectly affect the genes involved in invasion perhaps through the interaction between tumors and surrounding tissues. Because histological damage to the non-tumor regions was not observed after the administration of ARL4C ASO-1316(Harada et al, 2019), ARL4C might represent an appropriate target for pancreatic cancer therapy.

## Materials and Methods

### Materials and chemicals

HeLaS3 cells were kindly provided by Dr. K. Matsumoto (Nagoya University, Aichi, Japan) in May 2002. S2-CP8 pancreatic cancer cells were purchased from Cell Resource Center for Biomedical Research, Institute of Development, Aging and Cancer, Tohoku University, in April 2014. Lenti-X™ 293T (X293T) cells were purchased from Takara Bio Inc. (Shiga, Japan) in October 2011. PANC-1 cells were purchased from RIKEN Bioresource Center Cell Bank (Tsukuba, Japan) in October 2014. BxPC-3 cells were purchased from American Type Culture Collection in May 2018. HPAF-II cells were purchased from American Type Culture Collection in July 2017. S2-CP8, X293T, HeLaS3, Capan-2, Aspc-1, HPAF-II, and MDA-MB-231 cells were grown in Dulbecco’s modified Eagle’s medium (DMEM) supplemented with 10% fetal bovine serum (FBS). PANC-1 and BxPC-3 cells were grown in RPMI-1640 supplemented with 10% FBS.

S2-CP8 cells stably expressing GFP, ARL4C-EGFP, ARL4C^G2A^-EGFP, ARL4C^T27N^-EGFP, ARL4C^Q72L^-EGFP, ARL4C^ΔPBR^-EGFP, ARL4C^G2A^-EGFP-Cterm, ARL4C^ΔPBR^-EGFP-Cterm, ARL4C^G2A^-EGFP-GRP1^PH^, ARL4C^G2A^-EGFP-PLCδ^PH^, ARL4C-mCherry, ARL4C-tdTomato, EGFP-IQGAP1, and luciferase were generated using lentivirus as described previously (Kimura et al, 2016). BxPC-3 cells stably expressing EGFP or ARL4C-EGFP were generated using lentivirus. Lentiviral vector CSII-CMV-MCS-IRES2-Bsd harboring a cDNA was transfected with the packaging vectors pCAG-HIV-gp and pCMV-VSV-G-RSV-Rev into X293T cells using Lipofectamine2000 transfection reagent (Life Technologies/Thermo Fisher Scientific, Carlsbad, CA, USA). To generate S2-CP8 stable cells above, 1 x 10^5^ parental cells/well in a 12-well plate were treated with lentiviruses and 5 μg/mL polybrene, centrifuged at 1200 x *g* for 30 min, and incubated for 24 h. The cells were selected and maintained in the medium containing 10 μg/mL Blastcidin S.

ARL4C or IQGAP1 knockout cells were generated as previously described (Fujii et al, 2016). The target sequences for human ARL4C, 5’-CTTCTCGGTGTTGAAGCCGA-3’, and human IQGAP1, 5’-CACCGTGGGGTCTACCTTGCCAAAC-3’ were designed with the help of the CRISPR Genome Engineering Resources (http://www.genome-engineering.org/crispr/). The plasmids expressing hCas9 and single-guide RNA (sgRNA) were prepared by ligating oligonucleotides into the BbsI site of pX330 (addgene #42230). The plasmid pX330 with sgRNA sequences targeting ARL4C, IQGAP1 and Blasticidin resistance was introduced into S2-CP8 cells using Lipofectamine LTX reagent (Life Technologies/Thermo Fisher Scientific) according to manufacturer’s instructions and the transfected cells were selected in medium containing 5 μg/mL Blasticidin S for two days. Single colonies were picked, mechanically disaggregated, and replated into individual wells of 24-well plates.

ARL4C ASO-1316 and 6-carboxyfluorescein (FAM)-labelled ARL4C ASO-1316 were synthesized by GeneDesign (Osaka, Japan) as described(Harada et al, 2019). The sequences of the ASOs are listed in Supplementary file 1 Table 5. S2-CP8 cell were transfected with ASOs at 10 nmol/L using RNAiMAX (Life Technologies/Thermo Fisher Scientific) in antibiotics-free medium. The transfected cells were then used for experiments conducted at 48 h after transfection.

Anti-ARL4C polyclonal antibody (SAJ5550275) for immunoprecipitation and immunocytochemistry was generated in rabbits by immunization with recombinant human ARL4C. Antibodies used in this study are shown in Supplementary file 1 Table 6.

The following drugs were used: PD184161 (Sigma-Aldrich Co, St. Louis, MO, USA); U0126 (Promega Corp., Madison, WI, USA); Rapamycin (Cell Signaling Technology, Beverly, MA, USA); LY294002 (Cell Signaling Technology); and VivoGlo luciferin (Promega Corp.).

### Plasmid construction

pEGFPC2-IQGAP1, pEGFP-mCyth2, pAcGFP-mPlcd1^PH^, and CSII-CMV-MCS-IRES2-Bsd were kindly provided by K. Kaibuchi (Nagoya University, Japan), J. Yamauchi (Tokyo University of Pharmacy and Life Science, Japan), M. Matsuda (Kyoto University, Kyoto, Japan), and H. Miyoshi (RIKEN Bioresource Center, Ibaraki, Japan), respectively.

To generate plasmid DNA with mutated codons or deletions, site-directed mutagenesis method was performed using PrimeSTAR Max DNA Polymerase (Takara Bio Inc., Shiga, Japan). To generate plasmid DNA with insertions, PCR amplified fragments and linearized vector by restriction enzyme digestion were assembled using In-Fusion HD Cloning Kit (Takara Bio Inc.). pEGFPN3-ARL4C was constructed as previously described(Matsumoto et al, 2014). Full length cDNAs of GRP1 and MMP14 ORF were reversely transcribed from mRNA extracted from MCF-7 cells and U2OS cells, respectively. Linear double strand oligonucleotides of the C-terminal 22 amino acids of KRAS, which includes the PBR and CAAX motifs, were synthesized, and the oligonucleotides were inserted into C terminal of ARL4C-EGFP or ARL4C-FLAG-HA using In-Fusion HD Cloning Kit (Takara Bio Inc.).

Standard recombinant DNA techniques mentioned above were used to construct the following plasmids: pEGFPN3-ARL4C, pEGFPN3-ARL4C^G2A^, pEGFPN3-ARL4C^T27N^, pEGFPN3-ARL4C^Q72L^, pEGFPN3-ARL4C^ΔPBR^, pEGFPN3-ARL4C^G2A^-EGFP-PLCδ^PH^, pEGFPN3-ARL4C^G2A^-EGFP-GRP1^PH^, pEGFPN3-ARL4C^G2A^-EGFP-Cterm, pEGFPN3-ARL4CΔPBR-EGFP-Cterm, pEGFPC1-CHD, pEGFPC1-IQ, pEGFPC1-WW, pEGFPC1-IR, pEGFPC1-GRD, pEGFPC1-RGCT, pcDNA3-ARL4C-FLAG-HA, pcDNA3-ARL4C^G2A^-FLAG-HA, pcDNA3-ARL4C^ΔPBR^-FLAG-HA, pcDNA3-ARL4C^G2A^-FLAG-HA-Cterm, pcDNA3-ARL4C^ΔPBR^-FLAG-HA-Cterm, pcDNA3-FLAG-HA-IQGAP1, pmCherryN1-ARL4C, pmCherryN1-MMP14, pmCherryN1-MMP14ΔC(Δ563-582), pCAG-ARL4C-tdTomato. To construct lentiviral vectors harboring EGFP, ARL4C-EGFP, ARL4C^G2A^-EGFP, ARL4C^T27N^-EGFP, ARL4C^Q72L^-EGFP, ARL4C^ΔPBR^-EGFP, ARL4C^G2A^-EGFP-Cterm, ARL4C^ΔPBR^-EGFP-Cterm, ARL4C^G2A^-EGFP-PLCδPH, ARL4C^G2A^-EGFP-GRP1PH, EGFP-IQGAP1, ARLC-mCherry, MMP14-mCherry, MMP14ΔC-mCherry, ARL4C-tdTomato were cloned into CSII-CMV-MCS-IRES2-Bsd provided by Dr. H. Miyoshi (RIKEN Bioresource Center, Ibaraki, Japan).

### Patients and cancer tissues

The present study involved 57 presurgical untreated patients with PDAC and ages ranging from 47 to 87 years (median, 70 years) who underwent surgical resection at Osaka University between April 2001 and April 2015. Tumors were staged according to the Union for International Cancer Control (UICC) TNM staging system. Resected specimens were fixed in 10% (vol/vol) formalin, processed for paraffin embedding, and were sectioned at 5 μm thickness and stained with hematoxylin and eosin (H&E) or immunoperoxidase for independent evaluations. The protocol for this study was approved by the ethical review board of the Graduate School of Medicine, Osaka University, Japan (No. 13455), under the Declaration of Helsinki, and written informed consent was obtained from all patients. The study was performed in accordance with Committee guidelines and regulations.

### Immunohistochemical studies

Immunohistochemical studies were performed as previously described(Fujii et al, 2015) with modification. Briefly, all tissue sections were stained using a DakoReal EnVision Detection System (Dako, Carpentaria, CA, USA) in accordance with the manufacturer’s recommendations. Formalin-fixed, paraffin-embedded tissue specimens for examination were sectioned at 5-μm thickness. Heat-induced epitope retrieval was performed using Decloaking Chamber NxGen (Biocare Medical, Walnut Creek, CA, USA). Tissue peroxidase activity was blocked with Peroxidase-Blocking Solution (Dako) for 30 min, and the sections were then incubated with G-Block (GenoStaff, Tokyo, Japan) or Blocking One Histo (nacalai tesque, Kyoto, Japan) for 30 min or 10 min, respectively, to block nonspecific antibody binding sites. Tissue specimens were treated with anti-ARL4C (1:100), anti-IQGAP1 (1:800), or anti-MMP14 (1:100) antibody for 3 h at room temperature. Then, the specimens were detected by incubating with goat anti-rabbit or anti-rabbit/mouse IgG-HRP for 1 h and subsequently with DAB (Dako). The tissue sections were then counterstained with 0.1% (wt/vol) hematoxylin. ARL4C expression was considered high when the total area of the tumor stained with anti-ARL4C antibody exceeded 5%. IQGAP1 expression was considered high when the total area of the tumor stained with anti-IQGAP1 antibody exceeded 40%.

IQGAP1 staining positivity in PDAC patients was measured using HALO (Indica Labs, Corrales, NM, USA). The threshold for positive or negative staining was based on the optical density of the staining: regions above the positivity threshold were scored according to the optical density threshold set in the module; weakly positive is shown in yellow and strongly positive in red. The samples were viewed and analyzed using NanoZoomer-SQ (Hamamatsu Photonics K.K., Shizuoka, Japan).

### Clinical data analyses using open sources

The data on ARL4C and IQGAP1 mRNA expression in pancreatic adenocarcinoma were obtained from the UCSC Xena browser (http://xena.ucsc.edu). Tumors and normal samples in the UCSC Xena browser were derived from The Cancer Genome Atlas (TCGA) and Genotype-Tissue Expression (GTEx) projects. Differential analysis was performed using a two-tailed Student’s t-test. The correlations of overall survival rates with ARL4C, IQGAP1, and MMP14 expression in pancreatic cancer in TCGA datasets were analyzed using a Kaplan–Meier plotter (http://www.kmplot.com) and visualized using GraphPad Prism (GraphPad Software. San Diego, CA, USA). High and low expression groups were classified by auto select best cutoff. *P* values and *r* values were calculated using GraphPad Prism.

### 3D gel invasion assay using a 3D microfluidic cell culture chip

Collagen gels were made by diluting and neutralizing rat tail type I collagen (Corning Inc., Corning, NY, USA) in PBS and 12.1 mM NaOH, and was adjusted to 2 mg/mL. DQ™-collagen type I (Life Technologies/Thermo Fisher Scientific, Carlsbad, CA, USA) was mixed with collagen gels at a final concentration of 25 μg/mL. The gel channel of 3D microfluidic cell culture chip (AIM Biotech, Biopolis Rd, Singapore) was filled with collagen solution and incubated at 37°C for at least 1 h to polymerize collagen. After hydration of medium channels, a cell suspension (1 x 10^4^ cells) in serum-free cell culture medium with 0.2% BSA was injected into one of the ports at the medium channel. The opposite medium channel was filled with cell culture medium containing 10% FBS to create a chemoattractant gradient across the collagen gel. The cells were then incubated for 3 days and fixed for 15 min at room temperature in PBS containing 4% (w/v) paraformaldehyde. Then, the cells were permeabilized and blocked in PBS containing 0.5% (w/v) Triton X-100 and 40 mg/mL BSA for 30 min and stained with the indicated antibodies. The samples were viewed and analyzed under an LSM880 laser scanning microscope (Carl Zeiss, Jana, Germany). Reconstruction of confocal z-stack images into 3D animations and analysis of 4D images were performed using Imaris (Bitplane, Belfast, UK).

### Invadopodia assay

QCM™ Gelatin Invadopodia Assay (Red) (Merck Millipore, Burlington, MA, USA) was used in accordance with the manufacturer’s protocol. Briefly, poly-L-lysine–coated coverslips were treated with glutaraldehyde. The coverslips were then incubated with Cy3-labeled gelatin, followed by culture medium quenching of free aldehydes. Cells (6 x 10^4^ cells) were seeded onto the gelatin-coated coverslips and incubated for 4 h. After incubation, the cells were fixed for 20 min at room temperature in phosphate-buffered saline (PBS) containing 4% (w/v) paraformaldehyde and permeabilized in PBS containing 0.2% (w/v) Triton X-100 for 10 min. After being blocked in PBS containing 0.2% (w/v) BSA for 30 min, the cells were immunohistochemically stained. The samples were viewed and analyzed under an LSM880 laser scanning microscope (Carl Zeiss, Jana, Germany).

### 2D culture on poly-D-lysine– or Matrigel-coated dishes

Cells grown on glass coverslips coated with poly-D-lysine (Sigma-Aldrich) or Matrigel (Corning Inc., Corning, NY, USA) were fixed for 10 min at room temperature in PBS containing 4% (w/v) paraformaldehyde and permeabilized in PBS containing 0.1% (w/v) saponin (Sigma-Aldrich) or 0.2% (w/v) Triton X-100 for 10 min. The cells were then blocked in PBS containing 0.2% (w/v) BSA for 30 min. They were then incubated with primary antibodies for 3 h at room temperature and with secondary antibodies in accordance with the manufacturer’s protocol (Life Technologies/Thermo Fisher Scientific). For cell surface MMP14 staining, samples were incubated with anti-MMP14 antibody for 3 h at room temperature without permeabilization. The samples were viewed and analyzed under an LSM880 laser scanning microscope (Carl Zeiss).

### Migration and invasion assays

Migration and invasion assays were performed using a modified Boyden chamber (8 μm pores; Corning) and a Matrigel-coated modified Boyden chamber (8 μm pores; BD Biosciences, Franklin Lakes, NJ, USA), respectively as described previously(Kurayoshi et al, 2006; Matsumoto et al, 2014). In the standard conditions, S2-CP8 cells (2.5 x 10^4^ cells) were seeded in the upper side of Boyden Chamber. In GFP-expressing S2-CP8 cells, after 4 h (migration assay) or 24 h (invasion assay, except for Figure 6E) incubation with control ASO, 122 cells (average) and 126 cells (average), respectively, were observed in the lower side chamber in the one field of view under fluorescence microscope (BZ-9000, Keyence, Osaka, Japan) using a 10x air objective. In Figure 6E, cells were observed after 20 h incubation with ASO. Migration and invasion rates of cells expressing ARL4C, IQGAP1, and MMP14 mutants were calculated as the percentages of the same cells transfected with control ASO or siRNA.

### 3D type I collagen gel culture

Collagen gels were made by diluting and neutralizing rat tail type I collagen (Corning) in PBS and 12.1 mM NaOH, and was adjusted to 2 mg/mL. Then, 140 μL of cell-embedded collagen gels (1 x 10^6^ cells/mL) were overlaid onto glass coverslips in a 24-well plate and allowed to polymerize for at least 1 h at 37°C and 5% CO_2_. After polymerization, growth medium was added on top of the collagen gel. The cells were then incubated for 3 days and fixed for 15 min at room temperature in PBS containing 4% (w/v) paraformaldehyde. Then, the cells were permeabilized and blocked in PBS containing 0.5% (w/v) Triton X-100 and 40 mg/mL BSA for 30 min and incubated with primary antibodies for 3 h at room temperature and secondary antibodies in accordance with the manufacturer’s protocol (Life Technologies/Thermo Fisher Scientific). The samples were viewed and analyzed under an LSM880 laser scanning microscope using a 20x air objective (Carl Zeiss). In the standard conditions (for Figure 3L) with BxPC-3/ARL4C-GFP cells treated with control ASO, the number of cells with protrusions and the total number of cells were 15 (average) and 76 (average), respectively, in the one field of view under an LSM880 laser scanning microscope (Carl-Zeiss) using a 20x air objective. The percentages of cells with protrusions compared with the total number of cells in the presence of control siRNA or IQGAP1 siRNA were calculated.

### Inducible recruitment of phospholipid phosphatases

mRFP-FKBP-5-ptase-dom and PM-FRB-CFP plasmids were obtained from Addgene (deposited by the laboratory of T. Balla). S2-CP8 cells were then transiently transfected with both mRFP-FKBP-5-ptase-dom and PM-FRB-CFP (0.5 μg/well of a 6-well plate for each vector) with ViaFect (Promega Corp.). After 24 h culture, the cells were treated with 100 nM rapamycin or 50 μM LY294002 for 30 min before fixation.

### Isolation of ARL4C-interacting protein

Confluent X293T cells transiently transfected with ARL4C-FLAG-HA in two 10-cm culture dishes were harvested and lysed in 800 μL of lysis buffer (25mM Tris-HCl [pH7.5], 50 mM NaCl, 0.5% TritonX-100) with protease inhibitors (nacalai tesque). After 10 min of centrifugation, lysates were incubated with 40 μL of 50% slurry of anti-FLAG Affinity Gel for 30 min, and then add another 40 μL and incubated for 30 min. Beads were washed 3 times with 1 mL of lysis buffer. Recovered beads were incubated once with FLAG peptide (0.5 mg/mL) to elute proteins in 80 μL of PBS for 30 min at 4°C. Then, the supernatant was precleaned with 40 μL of 50% slurry of protein A Sepharose beads (GE Healthcare, Chicago, IL, USA) for 30 min at 4°C. The precleaned lysates were incubated with 2 μg of anti-HA antibody (Santa Cruz, Dallas, TX, USA) and 50 μL of 50% slurry of protein A Sepharose beads for 1 h at 4°C. Beads were washed 3 times with 1 mL of lysis buffer, and bound complexes were dissolved in 50 μL of Laemmli’s sample buffer. The ARL4C-FLAG-HA-interacting proteins were detected by silver staining (Life Technologies/Thermo Fisher Scientific). Nine bands (arrowheads in Figure 3A) were cut from the gel and analyzed by mass spectrometry.

### Immunoprecipitation

Immunoprecipitation were performed as described previously with modification(Matsumoto et al, 2014). For Figure 3C, S2-CP8 cells (60-mm diameter dish) were lysed in 300 μL of lysis buffer (25 mM Tris–HCl [pH 7.5, 50 mM NaCl, 0.5% Triton-X100) with protease inhibitors (nacalai tesque) for 10 min on ice. After centrifugation, the supernatant was collected and pre-cleaned using 30 μL of Dynabeads Protein G (Thermo Fisher Scientific). After pre-cleaning, lysates were rotated with complex of Dynabeads (50 μL) and antibody (3.6 μg) for 10 min at room temperature. The beads were then washed with lysis buffer three times, and finally suspended in Laemmli’s sample buffer.

### The RAC1 activity assay

The RAC1 activity assay was performed as described(Matsumoto et al, 2014). Briefly, cells were lysed in 400 μL of Rac1 assay buffer (20 mM Tris–HCl [pH 7.5], 150 mM NaCl, 1 mM dithiothreitol, 10 mM MgCl2, 1% Triton-X100) with protease inhibitors (nacalai tesque) containing 20 μg of glutathione-S-transferase (GST)-CRIB. After the lysates were centrifuged at 20,000 *g* for 10 min, the supernatants were incubated with glutathione-Sepharose (20 μl each) for 2 h at 4°C. The beads were then washed with Rac1 assay buffer three times, and finally suspended in Laemmli’s sample buffer. The precipitates were probed with the anti-Rac1 antibody.

### Imaging of ASO accumulation in tumor-bearing mice

Orthotopic transplantation was performed as described previously(Kim et al, 2009). Ten days after the transplantation, 150 μg/animal (approximately 7.5 mg/kg) of 6-FAM-ARL4CASO-1316 was subcutaneously administered. Four h after the injection, the fluorescence intensities of various organs were measured *ex vivo* using the IVIS imaging system (Xenogen Corp.). After *ex vivo* imaging, unfixed mouse pancreas tissues were frozen in an OCT (Sakura Finetek, Tokyo, Japan)/sucrose mixture [1:1 (v/v) OCT and 1 x PBS containing 30% sucrose]. Freshly frozen tissues were sectioned at 10 μm and fixed for 30 min at room temperature in PBS containing 4% (w/v) paraformaldehyde. The cells were then permeabilized and blocked in PBS containing 0.5% (w/v) Triton X-100 and 40 mg/mL BSA for 30 min and stained with the indicated antibodies. The samples were viewed and analyzed under an LSM880 laser scanning microscope (Carl Zeiss).

### Orthotopic xenograft tumor assay

An orthotopic transplantation assay was performed as described previously(Kim et al, 2009) with modification. Ten-week-old male BALB/cAJcl-nu/nu mice (nude mice; CLEA, Tokyo, Japan) were anesthetized and received an orthotopic injection of S2-CP8 cells into the mid-body of the pancreas using a 27 G needle (5 x 10^5^ cells suspended in 100 μL of HBSS with 50% Matrigel). ASOs (50 μg/mouse, approximately 2.5 mg/kg) were administered subcutaneously twice a week from day 3. Tumor burden was measured once a week using the IVIS imaging system (Xenogen Corp., Alameda, CA, USA). For the *in vivo* imaging, 100 μL of VivoGlo luciferin (30 mg/mL) was intraperitoneally administered and the bioluminescence imaging was performed 8 min later. The region of interest (ROI) was selected and the radiance values measured with Living Image 4.3.1 Software (Caliper Life Sciences, Hopkinton, MA, USA). The mice were euthanized 28 days after transplantation. Tumor weights and numbers of mesenteric lymph nodes (diameter of lymph nodes > 1 mm) were measured. All protocols used for the animal experiments in this study were approved by the Animal Research Committee of Osaka University, Japan (No. 26-032-048).

### RNA sequencing

Sequenced reads were preprocessed by Trim Galore! v0.6.3 and quantified by Salmon v0.14.0 with the flags gcBias and validateMappings. GENCODE vM21 annotation was used as the transcript reference. The quantified transcript-level scaled TPM was summarized into a gene-level scaled TPM by using the R package tximport v1.6.0. All procedures were implemented using the RNAseq pipeline ikra v1.2.2 [http://doi.org/10.5281/zenodo.3352573] with the default parameters. Downstream analysis was conducted with an integrative RNAseq analysis platform, iDEP.90. After normalization with VST, principal component analysis was conducted. Hierarchical clustering was performed on the top 1,000 genes in terms of their standard deviation. Finally, DEGs were selected with a log2 fold change > 1 and false discovery rate < 0.1.

### Ingenuity Pathway Analysis (IPA)

DEGs identified from RNA sequence data were subjected to Ingenuity Pathway Analysis (IPA; Qiagen, Hilden, Germany). This analysis examines DEGs that are known to affect each biological function and compares their direction of change to what is expected from the literature. To infer the activation states of implicated biological functions, two statistical quantities, Z-score and *P* value, were used. A positive or negative Z-score value indicates that biological functions are predicted to be activated or inhibited in the ARL4C ASO-1316–treated group relative to the control ASO-treated group. A negative Z-score means that the indicated biological functions are inhibited by ARL4C ASO-1316. The *P* value, calculated with the Fisher’s exact test, reflects the enrichment of the DEGs on each pathway. For stringent analysis, only biological functions with a |Z-score| > 2 were considered significant.

### Statistics and Reproducibility

Biological replicates are replicates on independent biological samples versus technical replicates that use the same starting samples. All experiments in this study were repeated using biological replicates. A minimum of three biological replicates were analyzed for all samples, and the results are presented as the mean ± s.d. or s.e.m. The cumulative probabilities of overall survival were determined using the Kaplan–Meier method; a log-rank test was used to assess statistical significance. The Student’s t-test or Mann-Whitney test was used to determine if there was a significant difference between the means of two groups. One-way analysis of variance (ANOVA) with Bonferroni tests was used to compare three or more group means. Statistical analysis was performed using Excel Toukei (ESUMI Co., Ltd., Tokyo, Japan); *P* < 0.05 was considered statistically significant. In box and whiskers plots, the top and bottom horizontal lines represent the 75^th^ and the 25^th^ percentiles, respectively, and the middle horizontal line represents the median. The size of the box represents the interquartile range and the top and bottom whiskers represent the maximum and the minimum values, respectively.

### Others

The siRNAs and primers used in these experiments are listed in Supplementary file 1 Table 7 and 8, respectively. 2.5D Matrigel growth assay and quantitative PCR were performed as described previously(Matsumoto et al, 2019; Sato et al, 2010).

## Acknowledgements

This work was supported by Grants-in-Aids for Scientific Research to A.K. (2016-2020) (No. 16H06374) and Grants-in-Aid for Scientific Research on Innovative Areas, “Organelle zone” to A.K. (2018-2019) (No. 18H04861) and “Cell diverse” to A.K. (2018-2019) (No. 18H05101) from the Ministry of Education, Culture, Sports, Science and Technology of Japan, by grants from the Yasuda Memorial Foundation and the Ichiro Kanehara Foundation of the Promotion of Medical Science & Medical Care to A.K., and by Integrated Frontier Research for Medical Science Division, Institute for Open and Transdisciplinary Research Initiatives, Osaka University to A.K.

We thank the NGS core facility of the Genome Information Research Center at the Research Institute for Microbial Diseases of Osaka University for the data analysis support.

This study was supported by Eiji Oiki, Yuri Terao, and Center for Medical Research and Education, Graduate School of Medicine, Osaka University. Also this study was supported by Saki Ishino and Center of Medical Innovation and Translational Research, Osaka University.

## Author contributions

Conceptualizaion: A.H., S.M. and A.K.; Methodology: A.H., S.M., T.A. and A.K.; Investigation: A.H., S.M. and Y.Y.; Resources: H.E.; Writing-original draft: A.H., S.M. and A.K.; Writing-review and editing: A.H., S.M., Y.Y., H.E. and A.K.; Supervision: A.K.; Project administration: A.H., S.M. and A.K.; Funding acquisition: A.K.

## Data availability

All data generated or analysed during this study are included in the manuscript and supporting files.

The following previously published data sets were used.

Normalized RNA-seq data and clinical information of pancreatic ductal adenocarcinoma samples from The Cancer Genome Atlas (TCGA) Research Network were downloaded from UCSC Xena website (https://xenabrowser.net/datapages/, 2020_04_07_run). Patients with missing or insufficient data were excluded from this research.

The following data sets were generated.

Akikazu Harada, Akira Kikuchi (2021)

ID DRA011537.

Effects of ARL4C ASO on an orthotopic transplantation model.

## Supplementary files

Figure 2-video 1; ARL4C accumulates at the tips of membrane protrusions.

Legend for Figure 2-video 1 is as follows.

S2-CP8 cells stably expressing ARL4C-tdTomato were subjected to the 3D gel invasion assay and were observed with time-lapse imaging and the video was acquired for 78 min. Cells were imaged every 3 min.

Figure 1-figure supplement 1

Figure 2-figure supplement 1

Figure 2-figure supplement 2

Figure 3-figure supplement 1

Figure 3-figure supplement 2

Figure 4-figure supplement 1

Figure 5-figure supplement 1

Figure 6-figure supplement 1

Figure 7-figure supplement 1

Figure 1-source data; Excel file containing quantitative data for Figure 1.

Figure 2-source data; Excel file containing quantitative data for Figure 2.

Figure 3-source data; Excel file containing quantitative data for Figure 3.

Figure 4-source data; Excel file containing quantitative data for Figure 4.

Figure 5-source data; Excel file containing quantitative data for Figure 5.

Figure 6-source data; Excel file containing quantitative data for Figure 6.

Figure 7-source data; Excel file containing quantitative data for Figure 7.

Figure 2-figure supplement 1-source data; Excel file containing quantitative data for Figure 2-figure supplement 1.

Figure 2-figure supplement 2-source data; Excel file containing quantitative data for Figure 2-figure supplement 2.

Figure 3-figure supplement 1-source data; Excel file containing quantitative data for Figure 3-figure supplement 1.

Figure 3-figure supplement 2-source data; Excel file containing quantitative data for Figure 3-figure supplement 2.

Figure 6-figure supplement 1-source data; Excel file containing quantitative data for Figure 6-figure supplement 1.

Figure 7-figure supplement 1-source data; Excel file containing quantitative data for Figure 7-figure supplement 1.

**Supplementary file 1 Tables 1-8**

## References

Briggs MW, Sacks DB (2003) IQGAP proteins are integral components of cytoskeletal regulation. EMBO Rep 4:571–574: 10.1038/sj.embor.embor867

Burd CG, Strochlic TI, Gangi Setty SR (2004) Arf-like GTPases: not so Arf-like after all. Trends Cell Biol 14:687–694: 10.1016/j.tcb.2004.10.004

Castellano E, Downward J (2011) RAS Interaction with PI3K: More Than Just Another Effector Pathway. Genes Cancer 2:261–274: 10.1177/1947601911408079

Caswell PT, Zech T (2018) Actin-Based Cell Protrusion in a 3D Matrix. Trends Cell Biol 28:823–834: 10.1016/j.tcb.2018.06.003

Collins MA, Bednar F, Zhang Y, Brisset JC, Galban S, Galban CJ, Rakshit S, Flannagan KS, Adsay NV, Pasca di Magliano M (2012) Oncogenic Kras is required for both the initiation and maintenance of pancreatic cancer in mice. J Clin Invest 122:639–653: 10.1172/JCI59227

Dalaka E, Kronenberg NM, Liehm P, Segall JE, Prystowsky MB, Gather MC (2020) Direct measurement of vertical forces shows correlation between mechanical activity and proteolytic ability of invadopodia. Sci Adv 6:eaax6912: 10.1126/sciadv.aax6912

Di Paolo G, De Camilli P (2006) Phosphoinositides in cell regulation and membrane dynamics. Nature 443:651–657: 10.1038/nature05185

Donaldson JG, Jackson CL (2011) ARF family G proteins and their regulators: roles in membrane transport, development and disease. Nat Rev Mol Cell Biol 12:362–375: 10.1038/nrm3117

Engel T, Lueken A, Bode G, Hobohm U, Lorkowski S, Schlueter B, Rust S, Cullen P, Pech M, Assmann G, Seedorf U (2004) ADP-ribosylation factor (ARF)-like 7 (ARL7) is induced by cholesterol loading and participates in apolipoprotein AI-dependent cholesterol export. FEBS Lett 566:241–246: 10.1016/j.febslet.2004.04.048

Farahat WA, Wood LB, Zervantonakis IK, Schor A, Ong S, Neal D, Kamm RD, Asada HH (2012) Ensemble analysis of angiogenic growth in three-dimensional microfluidic cell cultures. PLoS One 7:e37333: 10.1371/journal.pone.0037333

Fujii S, Matsumoto S, Nojima S, Morii E, Kikuchi A (2015) Arl4c expression in colorectal and lung cancers promotes tumorigenesis and may represent a novel therapeutic target. Oncogene 34:4834–4844: 10.1038/onc.2014.402

Fujii S, Shinjo K, Matsumoto S, Harada T, Nojima S, Sato S, Usami Y, Toyosawa S, Morii E, Kondo Y, Kikuchi A (2016) Epigenetic upregulation of ARL4C, due to DNA hypomethylation in the 3’-untranslated region, promotes tumorigenesis of lung squamous cell carcinoma. Oncotarget 7:81571–81587: 10.18632/oncotarget.13147

Hancock JF, Paterson H, Marshall CJ (1990) A polybasic domain or palmitoylation is required in addition to the CAAX motif to localize p21ras to the plasma membrane. Cell 63:133–139:

Harada T, Matsumoto S, Hirota S, Kimura H, Fujii S, Kasahara Y, Gon H, Yoshida T, Itoh T, Haraguchi N, Mizushima T, Noda T, Eguchi H, Nojima S, Morii E, Fukumoto T, Obika S, Kikuchi A (2019) Chemically modified antisense oligonucleotide against ARL4C inhibits primary and metastatic liver tumor growth. Mol Cancer Ther 18:602–612: 10.1158/1535-7163.MCT-18-0824

Hedman AC, Smith JM, Sacks DB (2015) The biology of IQGAP proteins: beyond the cytoskeleton. EMBO Rep 16:427–446: 10.15252/embr.201439834

Heo WD, Inoue T, Park WS, Kim ML, Park BO, Wandless TJ, Meyer T (2006) PI(3,4,5)P3 and PI(4,5)P2 lipids target proteins with polybasic clusters to the plasma membrane. Science 314:1458–1461: 10.1126/science.1134389

Hidalgo M (2010) Pancreatic cancer. N Engl J Med 362:1605–1617: 10.1056/NEJMra0901557

Hofmann I, Thompson A, Sanderson CM, Munro S (2007) The Arl4 family of small G proteins can recruit the cytohesin Arf6 exchange factors to the plasma membrane. Curr Biol 17:711–716: 10.1016/j.cub.2007.03.007

Jacquemet G, Green DM, Bridgewater RE, von Kriegsheim A, Humphries MJ, Norman JC, Caswell PT (2013) RCP-driven α5β1 recycling suppresses Rac and promotes RhoA activity via the RacGAP1-IQGAP1 complex. J Cell Biol 202:917–935: 10.1083/jcb.201302041

Jiang A, Lehti K, Wang X, Weiss SJ, Keski-Oja J, Pei D (2001) Regulation of membrane-type matrix metalloproteinase 1 activity by dynamin-mediated endocytosis. Proc Natl Acad Sci USA 98:13693–13698: 10.1073/pnas.241293698

Johnson M, Sharma M, Henderson BR (2009) IQGAP1 regulation and roles in cancer. Cell Signal 21:1471–1478: 10.1016/j.cellsig.2009.02.023

Keleg S, Buchler P, Ludwig R, Buchler MW, Friess H (2003) Invasion and metastasis in pancreatic cancer. Mol Cancer 2:14:

Kim MP, Evans DB, Wang H, Abbruzzese JL, Fleming JB, Gallick GE (2009) Generation of orthotopic and heterotopic human pancreatic cancer xenografts in immunodeficient mice. Nat Protoc 4:1670–1680: 10.1038/nprot.2009.171

Kimura H, Fumoto K, Shojima K, Nojima S, Osugi Y, Tomihara H, Eguchi H, Shintani Y, Endo H, Inoue M, Doki Y, Okumura M, Morii E, Kikuchi A (2016) CKAP4 is a Dickkopf1 receptor and is involved in tumor progression. J Clin Invest 126:2689–2705: 10.1172/JCI84658

Kimura K, Matsumoto S, Harada T, Morii E, Nagatomo I, Shintani Y, Kikuchi A (2020) ARL4C is associated with initiation and progression of lung adenocarcinoma and represents a therapeutic target. Cancer Sci 111:951–961: 10.1111/cas.14303

Klein AP (2013) Identifying people at a high risk of developing pancreatic cancer. Nat Rev Cancer 13:66–74: 10.1038/nrc3420

Kurayoshi M, Oue N, Yamamoto H, Kishida M, Inoue A, Asahara T, Yasui W, Kikuchi A (2006) Expression of Wnt-5a is correlated with aggressiveness of gastric cancer by stimulating cell migration and invasion. Cancer Res 66:10439–10448: 10.1158/0008-5472.CAN-06-2359

Lemmon MA (2008) Membrane recognition by phospholipid-binding domains. Nat Rev Mol Cell Biol 9:99–111: 10.1038/nrm2328

Liang D, Shi S, Xu J, Zhang B, Qin Y, Ji S, Xu W, Liu J, Liu L, Liu C, Long J, Ni Q, Yu X (2016) New insights into perineural invasion of pancreatic cancer: More than pain. Biochim Biophys Acta 1865:111–122: 10.1016/j.bbcan.2016.01.002

Maffucci T, Falasca M (2001) Specificity in pleckstrin homology (PH) domain membrane targeting: a role for a phosphoinositide-protein co-operative mechanism. FEBS Let**t** 506:173–179: 10.1016/s0014-5793(01)02909-x

Matsumoto S, Fujii S, Kikuchi A (2017) Arl4c is a key regulator of tubulogenesis and tumourigenesis as a target gene of Wnt-β-catenin and growth factor-Ras signalling. J Biochem 161:27–35: 10.1093/jb/mvw069

Matsumoto S, Fujii S, Sato A, Ibuka S, Kagawa Y, Ishii M, Kikuchi A (2014) A combination of Wnt and growth factor signaling induces Arl4c expression to form epithelial tubular structures. EMBO J 33:702–718: 10.1002/embj.201386942

Matsumoto S, Yamamichi T, Shinzawa K, Kasahara Y, Nojima S, Kodama T, Obika S, Takehara T, Morii E, Okuyama H, Kikuchi A (2019) GREB1 induced by Wnt signaling promotes development of hepatoblastoma by suppressing TGFβ signaling. Nat Commun 10:3882: 10.1038/s41467-019-11533-x

Murphy DA, Courtneidge SA (2011) The ‘ins’ and ‘outs’ of podosomes and invadopodia: characteristics, formation and function. Nat Rev Mol Cell Biol 12:413–426: 10.1038/nrm3141

Pasqualato S, Renault L, Cherfils J (2002) Arf, Arl, Arp and Sar proteins: a family of GTP-binding proteins with a structural device for ‘front-back’ communication. EMBO Rep 3:1035–1041: 10.1093/embo-reports/kvf221

Petrie RJ, Gavara N, Chadwick RS, Yamada KM (2012) Nonpolarized signaling reveals two distinct modes of 3D cell migration. J Cell Biol 197:439–455: 10.1083/jcb.201201124

Poincloux R, Lizarraga F, Chavrier P (2009) Matrix invasion by tumour cells: a focus on MT1-MMP trafficking to invadopodia. J Cell Sci 122:3015–3024: 10.1242/jcs.034561

Rodriguez-Viciana P, Warne PH, Dhand R, Vanhaesebroeck B, Gout I, Fry MJ, Waterfield MD, Downward J (1994) Phosphatidylinositol-3-OH kinase as a direct target of Ras. Nature 370:527–532:

Sakurai-Yageta M, Recchi C, Le Dez G, Sibarita JB, Daviet L, Camonis J, D’Souza-Schorey C, Chavrier P (2008) The interaction of IQGAP1 with the exocyst complex is required for tumor cell invasion downstream of Cdc42 and RhoA. J Cell Biol 181:985–998: 10.1083/jcb.200709076

Sato A, Yamamoto H, Sakane H, Koyama H, Kikuchi A (2010) Wnt5a regulates distinct signalling pathways by binding to Frizzled2. EMBO J 29:41–54: emboj2009322 [pii] 10.1038/emboj.2009.322

Saykali BA, El-Sibai M (2014) Invadopodia, regulation, and assembly in cancer cell invasion. Cell Commun Adhes 21:207–212: 10.3109/15419061.2014.923845

Shin Y, Han S, Jeon JS, Yamamoto K, Zervantonakis IK, Sudo R, Kamm RD, Chung S (2012) Microfluidic assay for simultaneous culture of multiple cell types on surfaces or within hydrogels. Nat Protoc 7:1247–1259: 10.1038/nprot.2012.051

Suh BC, Inoue T, Meyer T, Hille B (2006) Rapid chemically induced changes of PtdIns(4,5)P2 gate KCNQ ion channels. Science 314:1454–1457: 10.1126/science.1131163

Toker A, Cantley LC (1997) Signalling through the lipid products of phosphoinositide-3-OH kinase. Nature 387:673–676: 10.1038/42648

Waddell N, Pajic M, Patch AM, Chang DK, Kassahn KS, Bailey P, Johns AL, Miller D, Nones K, Quek K, Quinn MC, Robertson AJ, Fadlullah MZ, Bruxner TJ, Christ AN, Harliwong I, Idrisoglu S, Manning S, Nourse C, Nourbakhsh E, Wani S, Wilson PJ, Markham E, Cloonan N, Anderson MJ, Fink JL, Holmes O, Kazakoff SH, Leonard C, Newell F, Poudel B, Song S, Taylor D, Waddell N, Wood S, Xu Q, Wu J, Pinese M, Cowley MJ, Lee HC, Jones MD, Nagrial AM, Humphris J, Chantrill LA, Chin V, Steinmann AM, Mawson A, Humphrey ES, Colvin EK, Chou A, Scarlett CJ, Pinho AV, Giry-Laterriere M, Rooman I, Samra JS, Kench JG, Pettitt JA, Merrett ND, Toon C, Epari K, Nguyen NQ, Barbour A, Zeps N, Jamieson NB, Graham JS, Niclou SP, Bjerkvig R, Grutzmann R, Aust D, Hruban RH, Maitra A, Iacobuzio-Donahue CA, Wolfgang CL, Morgan RA, Lawlor RT, Corbo V, Bassi C, Falconi M, Zamboni G, Tortora G, Tempero MA, Australian Pancreatic Cancer Genome I, Gill AJ, Eshleman JR, Pilarsky C, Scarpa A, Musgrove EA, Pearson JV, Biankin AV, Grimmond SM (2015) Whole genomes redefine the mutational landscape of pancreatic cancer. Nature 518:495–501: 10.1038/nature14169

Wei SM, Xie CG, Abe Y, Cai JT (2009) ADP-ribosylation factor like 7 (ARL7) interacts with alpha-tubulin and modulates intracellular vesicular transport. Biochem Biophys Res Commun 384:352–356: 10.1016/j.bbrc.2009.04.125

Wolf K, Wu YI, Liu Y, Geiger J, Tam E, Overall C, Stack MS, Friedl P (2007) Multi-step pericellular proteolysis controls the transition from individual to collective cancer cell invasion. Nat Cell Biol 9:893–904: 10.1038/ncb1616

