## Supplementary files for "Recruitment of KRAS downstream target ARL4C to membrane protrusions accelerates pancreatic cancer cell invasion"

Figure supplements and associated legends.
Supplementary file 1 Tables 1-8.

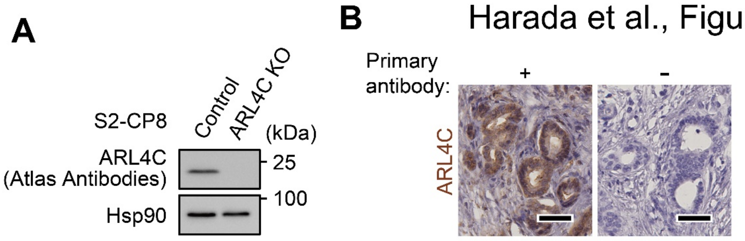


**Figure 1-figure supplement 1.**
**ARL4C is expressed in pancreatic cancer cells.**
**A,** Lysates were prepared from S2-CP8 WT or ARL4C KO cells and probed with the indicated antibodies. **B,** PDAC tissues were stained with or without anti-ARL4C antibody as the primary antibody and hematoxylin. Scale bars in **B,** 50 µm. KO, knockout.

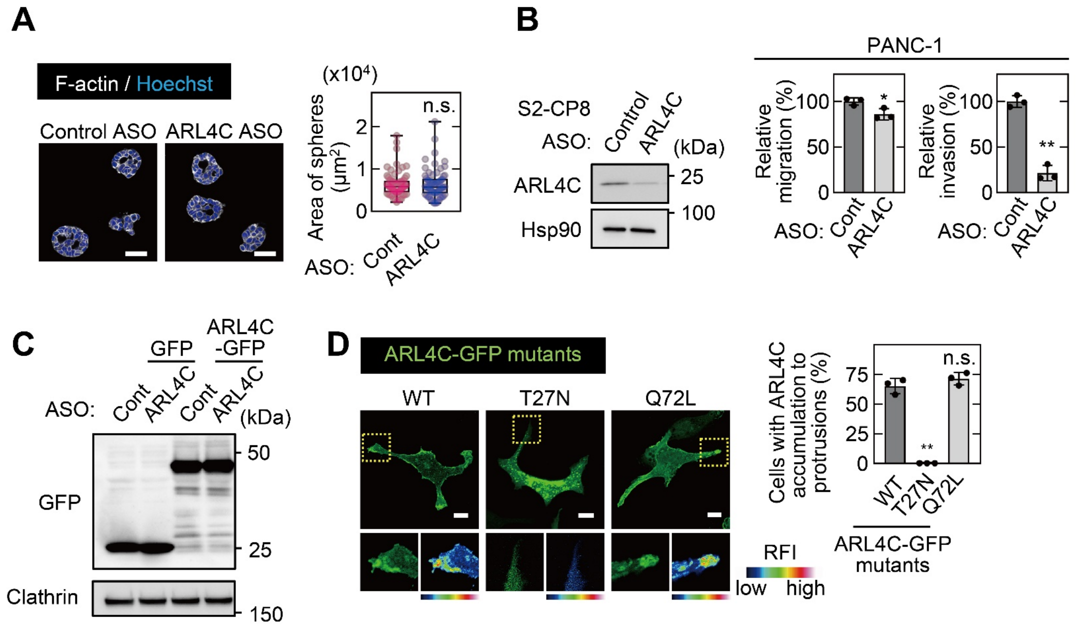


**Figure 2-figure supplement 1.**
**ARL4C expression is involved in invasion of pancreatic cancer cells rather than in sphere formation.**
**A,** S2-CP8 cells transfected with control or ARL4C ASO-1316 were cultured for 6 days in 2.5D Matrigel. The cells were then fixed and stained with phalloidin and Hoechst 33342 and sphere areas were calculated. When more than 10 cells formed a spherical structure, it was counted as one sphere. Data are shown as a box and whiskers plot. Center lines show the medians; box limits indicate the 25$^{th}$ and 75$^{th}$ percentiles; whiskers indicate the smallest and largest values; dots show all of the individual values. More than 65 spheres were analyzed per group. *P* values were calculated using a two-tailed Student's t-test. **B,** Lysates were prepared from S2-CP8 cells transfected with control or ARL4C ASO-1316 and probed with the indicated antibodies. PANC-1 cells transfected with control or ARL4C ASOs were subjected to migration and invasion assays. Migratory and invasive activities are expressed as the percentage of control cells. **C,** S2-CP8 cells stably expressing GFP or ARL4C-GFP were transfected with control or ARL4C ASO-1316. Lysates were probed with the indicated antibodies. **D,** S2-CP8 cells were transfected with the indicated ARL4C-GFP mutants. The regions in the yellow dashed squares are shown enlarged in the left bottom images. The right bottom images are shown in a false color representation of fluorescence intensity. The percentages of cells with ARL4C-GFP mutant accumulated at membrane protrusions compared with the total number of cells were calculated. False color representations were color-coded on the spectrum. **B,D,** Data are shown as the mean ± s.d. of 3 independent experiments. *P* values were calculated using a two-tailed Student's t-test (**B**) or one-way ANOVA followed by Bonferroni post hoc test (**D**). Scale bars in **A,** 50 µm; **D,** 10 µm. RFI, relative fluorescence intensity. n.s., not significant. *, $P < 0.05$; **, $P < 0.01$. See Figure 2-figure supplement 1-source data.

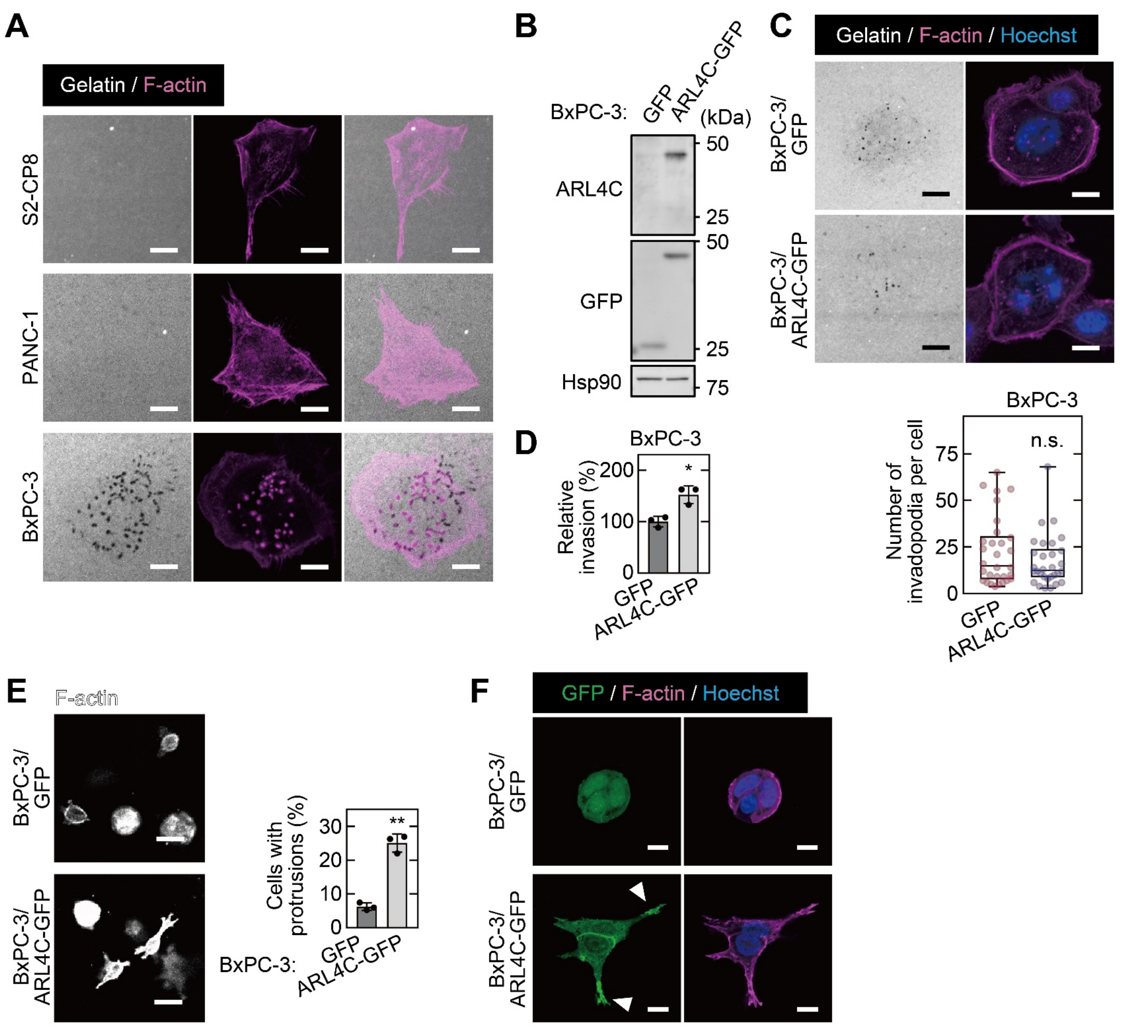


**Figure 2-figure supplement 2.**
**ARL4C is not involved in invadopodia formation.**
**A,** S2-CP8, PANC-1, and BxPC-3 cells were subjected to an invadopodia assay and then stained with phalloidin. **B,** Lysates of BxPC-3 cells expressing GFP or ARL4C-GFP were probed with the indicated antibodies. **C,** BxPC-3 cells expressing GFP or ARL4C-GFP were subjected to an invadopodia assay and stained with phalloidin and Hoechst 33342. Numbers of invadopodia per cell were counted. **D,** BxPC-3 cells expressing GFP or ARL4C-GFP were subjected to an invasion assay. Invasive activities are expressed as the percentage of control cells. **E,F,** BxPC-3 cells expressing GFP or ARL4C-GFP were cultured under 3D conditions with 2 mg/mL type I collagen for 3 days and stained with phalloidin and Hoechst 33342. Percentages of cells with cell protrusions compared with the total number of cells were calculated. White arrowheads in (**F**) indicate ARL4C-GFP accumulation at the cell protrusion. **D,E,** Data are shown as the mean ± s.d. of 3 independent experiments. *P* values were calculated using a two-tailed Student's t-test. **C,** Data are shown as a box and whiskers plot. Center lines show the medians; box limits indicate the 25th and 75th percentiles; whiskers indicate the smallest and largest values; dots show all of the individual values. 31 cells were analyzed per group. *P* values were calculated using the Mann–Whitney test. Scale bars in **A,C,F,** 10 µm; **E,** 50 µm. n.s., not significant. *, $P < 0.05$; **, $P < 0.01$. See Figure 2-figure supplement 2-source data.

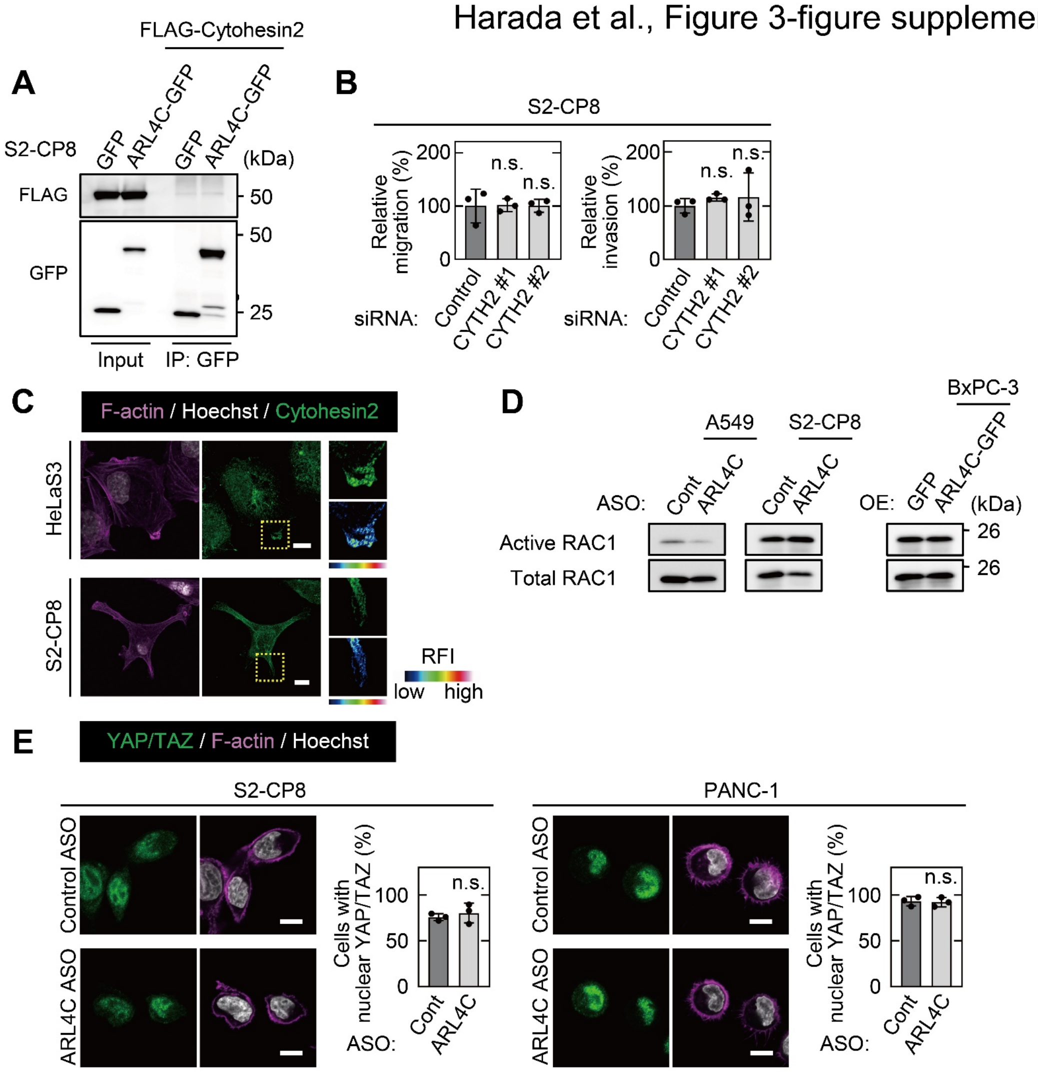


**Figure 3-figure supplement 1.**
**Cytohesin2 does not mediate ARL4C signaling in pancreatic cancer cells.**
**A,** FLAG-cytohesin2 was expressed in S2-CP8 cells expressing GFP or ARL4C-GFP. Lysates were immunoprecipitated with anti-GFP antibody, and the immunoprecipitates were probed with the indicated antibodies. **B,** S2-CP8 cells transfected with control or two independent CYTH2 (a gene of cytohesin2) siRNAs were subjected to migration and invasion assays. Migratory and invasive activities are expressed as the percentage of control cells. **C,** HeLaS3 and S2-CP8 cells were stained with anti-Cytohesin2 antibody, phalloidin, and Hoechst 33342. Enlarged images of the regions in the yellow dashed squares are shown in a false color representation of fluorescence intensity on the right. **D,** A549 and S2-CP8 cells transfected with the indicated ASOs, and BxPC-3 cells expressing GFP or ARL4C-GFP were subjected to assay for RAC1 activity. **E,** S2-CP8 and PANC-1 cells transfected with the indicated ASOs were cultured for 2.5 h under 2.5D Matrigel conditions and stained with anti-YAP/TAZ antibody and Hoechst 33342. Cells with nuclear YAP/TAZ were counted, and the data are shown as the percentage of positively stained cells compared with the total number of Hoechst-stained cells. **C,** False color representations were color-coded on the spectrum. **B,E,** Data are shown as the mean ± s.d. of 3 independent experiments. *P* values were calculated using a two-tailed Student's t-test (**E**) or one-way ANOVA followed by Bonferroni post hoc test (**B**). Scale bars in **C,E,** 10 µm. OE, overexpression. RFI, relative fluorescence intensity. n.s., not significant. See Figure 3-figure supplement 1-source data.

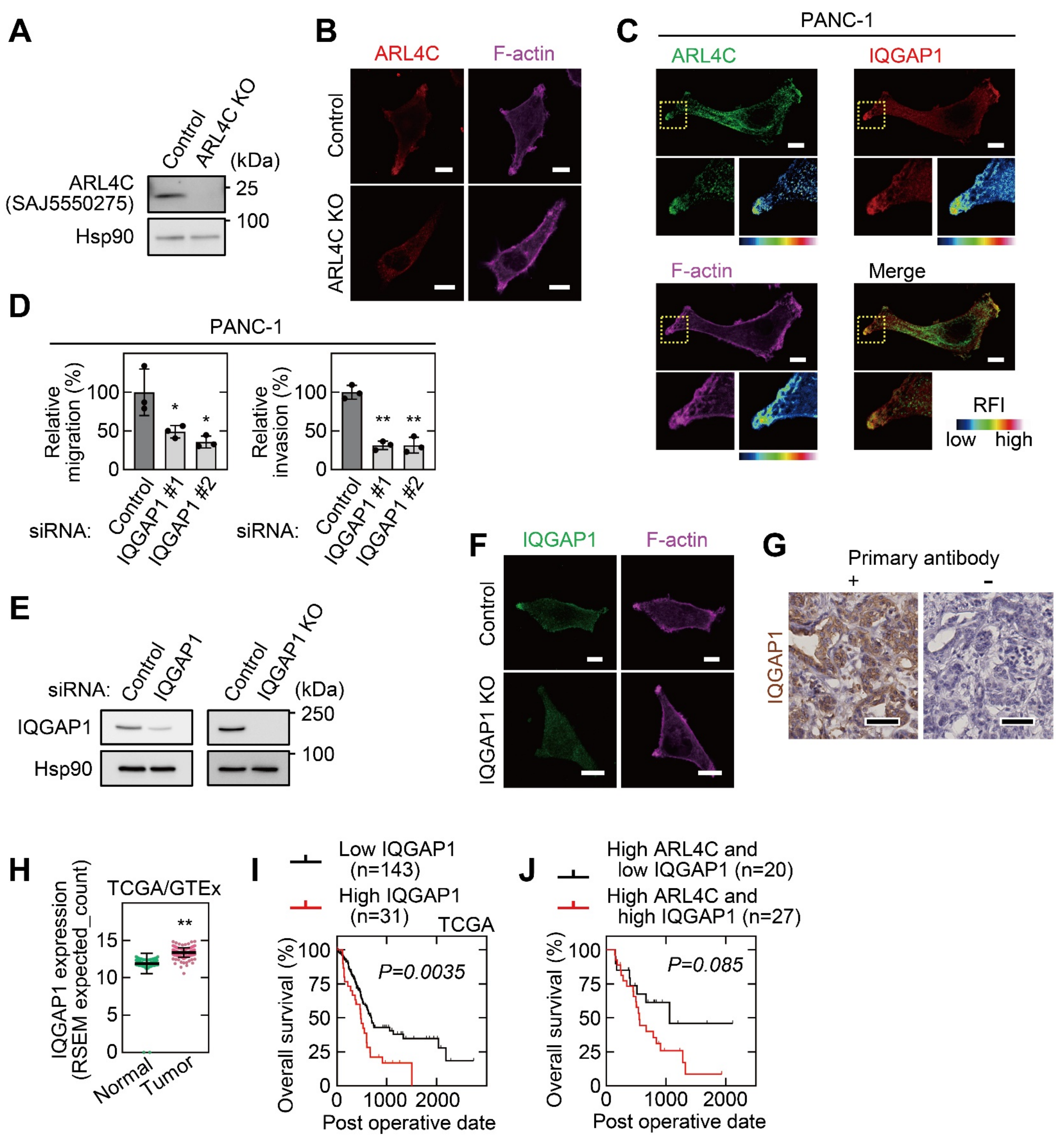


**Figure 3-figure supplement 2.**
**IQGAP1 interacts with ARL4C and involves in the invasion of pancreatic cancer cells.**
**A,** Lysates from S2-CP8 WT or ARL4C KO cells were probed with the indicated antibodies. **B,** S2-CP8 WT or ARL4C KO cells were stained with anti-ARL4C antibody and phalloidin. **C,** PANC-1 cells were stained with the indicated antibodies. Images of ARL4C and IQGAP1 were merged. Enlarged images of the regions in the yellow dashed squares are shown in a false color representation of fluorescence intensity on the bottom right. False color representations were color-coded on the spectrum. **D,** PANC-1 cells transfected with control or IQGAP1 siRNAs were subjected to migration and invasion assays. Migratory and invasive activities are expressed as the percentage of control cells. Data are shown as the mean ± s.d. of 3 independent experiments. *P* values were calculated using one-way ANOVA followed by Bonferroni post hoc test. **E,** Lysates were prepared from S2-CP8 cells transfected with control or IQGAP1 siRNA, and S2-CP8 WT or IQGAP1 KO cells. Lysates were probed with the indicated antibodies. **F,** S2-CP8 WT or IQGAP1 KO cells were stained with anti-IQGAP1 antibody and phalloidin. **G,** PDAC tissues were stained with or without anti-IQGAP1 antibody and hematoxylin. **H,** *IQGAP1* mRNA levels in pancreatic adenocarcinoma and normal tissues of the pancreas were analyzed using TCGA and GTEx datasets. The results are shown as scatter plots with the mean ± s.e.m. *P* values were calculated using a two-tailed Student's t-test. **I,** TCGA RNA sequencing and clinical outcome data for pancreatic cancer were analyzed. **J**, The relationship between overall survival and IQGAP1 expression in PDAC patients with high ARL4C expression was analyzed. **I,J,** The data were analyzed by Kaplan–Meier survival curves, and a log-rank test was used for statistical analysis. Scale bars in **B,C,F,** 10 µm; **G,** 50 µm. KO, knockout. RFI, relative fluorescence intensity. *, $P < 0.05$; **, $P < 0.01$. See Figure 3-figure supplement 2-source data.

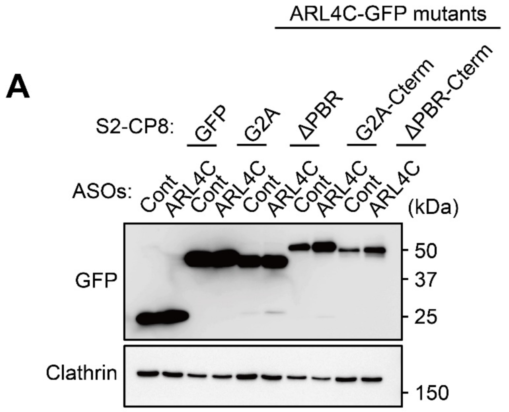


**Figure 4-figure supplement 1.**
**Expression of ARL4C mutants.**
**A,** S2-CP8 cells expressing GFP or the indicated mutants of ARL4C-GFP were transfected with control or ARL4C ASO-1316. Lysates were probed with the indicated antibodies.

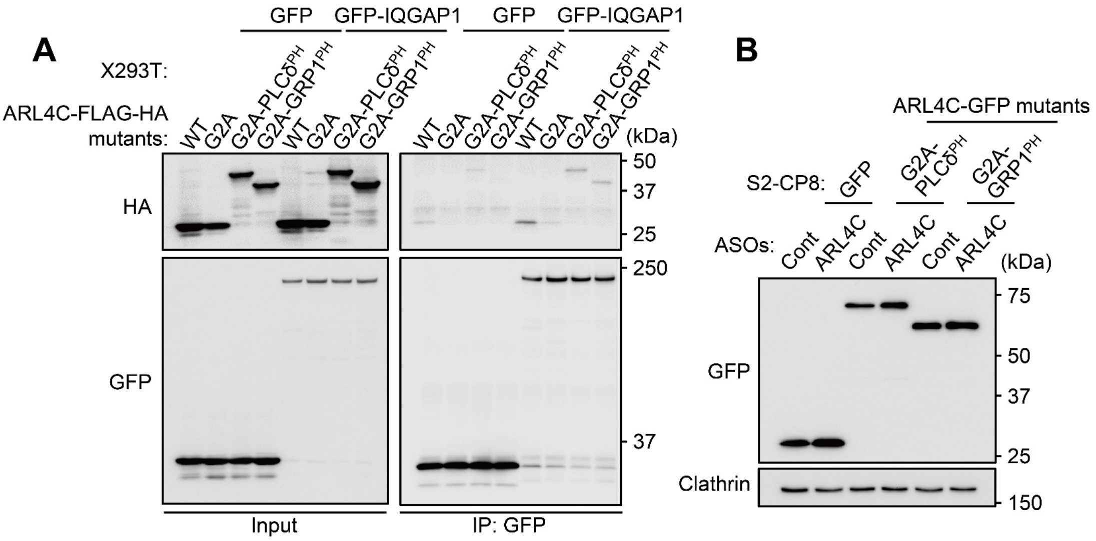


**Figure 5-figure supplement 1.**
**Interaction of ARL4C mutants and IQGAP1.**
**A,** Lysates of X293T cells expressing the indicated mutants of ARL4C-FLAG-HA and GFP or GFP-IQGAP1 proteins were immunoprecipitated with anti-GFP antibody, and the immunoprecipitates were probed with the indicated antibodies. **B,** S2-CP8 cells expressing GFP or the indicated mutants of ARL4C-GFP were transfected with control or ARL4C ASO-1316. Lysates were probed with the indicated antibodies.

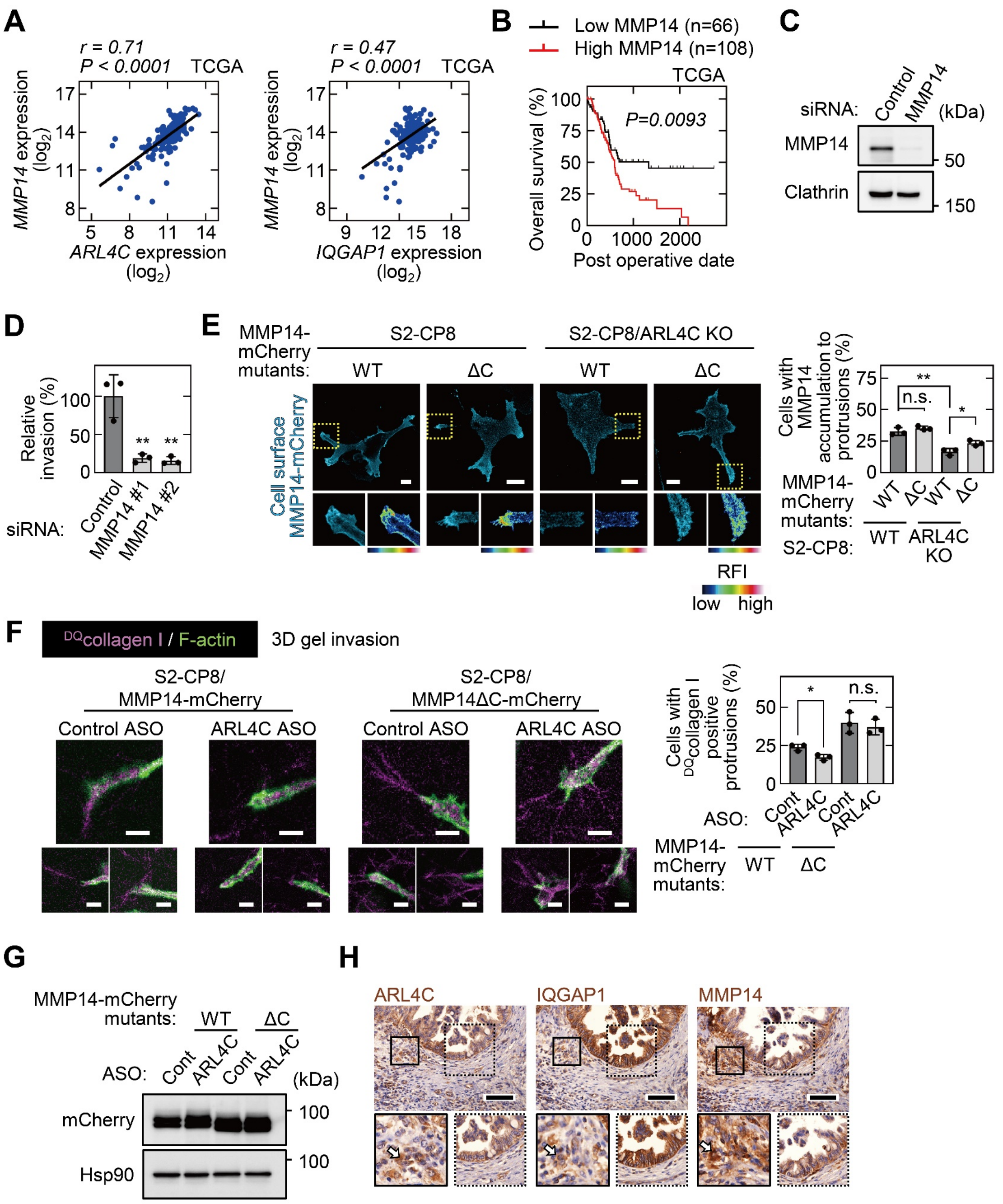

**Figure 6-figure supplement 1.**
**ARL4C recruits MMP14 to membrane protrusions and their expression is associated with poor prognosis in pancreatic cancer patients.** **A,** Scatter plot showing the correlation between the mRNA expression levels of *ARL4C* or *IQGAP1* (X-axis) and *MMP14* (Y-axis) in pancreatic cancer patients obtained from TCGA datasets using the R2: Genomics Analysis and Visualization Platform. *r* indicates the Pearson's correlation coefficient. **B,** TCGA RNA sequencing and clinical outcome data for pancreatic cancer were analyzed. The data were analyzed by Kaplan–Meier survival curves, and a log-rank test was used for statistical analysis. **C,** Lysates of S2-CP8 cells transfected with control or MMP14 siRNA were probed with anti-MMP14 and anti-Clathrin antibodies. **D,** S2-CP8 cells transfected with control or MMP14 siRNAs were subjected to invasion assay. Invasive activities are expressed as the percentage of control cells. **E,** S2-CP8 cells or ARL4C KO S2-CP8 cells expressing MMP14-mCherry or MMP14ΔC-mCherry were probed with anti-MMP14 antibody without permeabilization. The regions in the yellow dashed squares are shown enlarged in the left bottom images. The right bottom images are shown in a false color representation of fluorescence intensity. False color representations were color-coded on the spectrum. The percentages of cells with MMP14 accumulated at membrane protrusions compared with the total number of cells were calculated. **F,** S2-CP8 cells stably expressing MMP14-mCherry and MMP14ΔC-mCherry were transfected with control or ARL4C ASO-1316 and were then subjected to a 3D collagen I gel invasion assay with $^{DQ}$collagen I. The cells were stained with phalloidin. Three representative images for each condition are shown. Percentages of cells with $^{DQ}$collagen I-positive protrusions compared with the total number of cells were calculated. **G,** S2-CP8 cells stably expressing MMP14-mCherry and MMP14ΔC-mCherry were transfected with control or ARL4C ASO-1316. Lysates were probed with the indicated antibodies. **H,** PDAC tissues were stained with the indicated antibodies and hematoxylin. Magnified images are shown below. White arrows indicate cells invading into the surrounding interstitial tissues. 9 patient samples were imaged and the representative images are shown. **D-F,** Data are shown as the mean ± s.d. of 3 independent experiments. *P* values were calculated using a two-tailed Student's t-test (**F**) or one-way ANOVA followed by Bonferroni post hoc test (**D,E**). Scale bars in **E,F,** 10 μm; **H,** 100 μm. KO, knockout. RFI, relative fluorescence intensity. n.s., not significant. *, $P < 0.05$; **, $P < 0.01$. See Figure 6-figure supplement 1-source data.

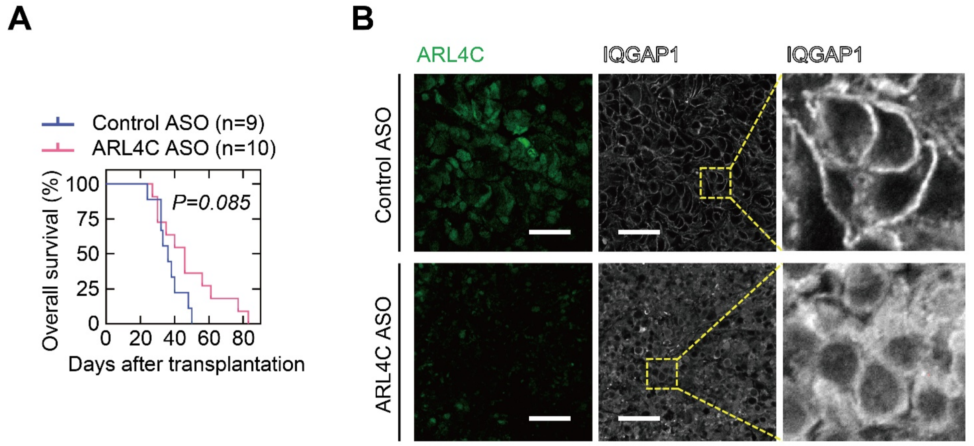


**Figure 7-figure supplement 1.**
**ARL4C ASO-1316 extends the survival of orthotopically transplanted mice.**
**A,** S2-CP8/Luciferase cells were implanted into the pancreas of nude mice, and control ASO (n = 9) or ARL4C ASO-1316 (n = 10) were administered subcutaneously twice a week. The Kaplan–Meier survival curve for the mice is shown. Statistical significance was determined by a log-rank test. **B,** PDAC tissues were stained with the indicated antibodies. Panels on the right show enlarged images of the yellow dashed squares. Scale bar in **B,** 50 µm. See Figure 7-figure supplement 1-source data.

**Supplementary File 1 Table 1.** Clinicopathological features of high and low ARL4C expression in PDAC.

| **Parameters** | | **High ARL4C cases** | **Low ARL4C cases** | ***P* value** |
|---|---|---|---|---|
| Stage | IA-IIA | 20(76.9%) | 6 | 0.49 |
| | IIB-III | 27(87.1%) | 4 | |
| T classification | T1/2 | 6(66.7%) | 3 | 0.18 |
| | T3 | 41(85.4%) | 7 | |
| N classification | N0 | 20(76.9%) | 6 | 0.49 |
| | N1 | 27(87.1%) | 4 | |
| Lymphatic vessel invasion | ly0 | 13(72.2%) | 5 | 0.26 |
| | ly1/2 | 34(87.2%) | 5 | |
| Venous invasion | v0 | 28(82.4%) | 6 | 1 |
| | v1/2/3 | 19(82.6%) | 4 | |
| Perineural invasion | ne0 | 2(40.0%) | 3 | 0.033 |
| | ne1/2/3 | 45(86.5%) | 7 | |
| Age | <65 | 12(75.0%) | 4 | 0.44 |
| | ≧65 | 35(85.4%) | 6 | |
| Sex | Male | 24(80.0%) | 6 | 0.73 |
| | Female | 23(85.2%) | 4 | |
| Tumor location | Head | 32(84.2%) | 6 | 0.72 |
| | Body and/or Tail | 15(78.9%) | 4 | |

Associations between ARL4C expression and the clinicopathologic characteristics were investigated. *P* values were calculated using the Chi-square test. T1, tumor limited to the pancreas, 2 cm or less in greatest dimension. T2, tumor limited to the pancreas, more than 2 cm in greatest dimension. T3, tumor extends beyond the pancreas but without involvement of the celiac axis or the superior mesenteric artery. N0, no regional lymph node metastasis. N1, regional lymph node metastasis. ly0, no lymphatic vessel invasion. ly1, mild lymphatic vessel invasion. ly2, moderate lymphatic vessel invasion. v0, no venous invasion. v1, mild venous invasion. v2, moderate venous invasion. v3, severe venous invasion. ne0, no perineural invasion. ne1, mild perineural invasion. ne2, moderate perineural invasion. ne3, severe perineural invasion.

**Supplementary File 1 Table 2.** The list of ARL4C binding proteins

| Band number | Protein name |
|---|---|
| 1 | PRKDC |
| 2 | IQGAP1 |
| 3 | IARS |
| 4 | SMC3 |
| 5 | DHX9 |
| 6 | PARP1 |
| 7 | MATR3 |
| 8 | HNRNPU |
| 9 | NCL |

The list of ARL4C binding proteins was shown.

**Supplementary File 1 Table 3.** Clinicopathological features of high and low IQGAP1 expression in PDAC.

| Parameters | | High IQGAP1 cases | Low IQGAP1 cases | *P* value |
|---|---|---|---|---|
| Stage | IA-IIA | 14(53.8%) | 12 | 1 |
| | IIB-III | 17(54.8%) | 14 | |
| T classification | T1/2 | 5(55.6%) | 4 | 1 |
| | T3 | 26(54.2%) | 22 | |
| N classification | N0 | 14(53.8%) | 12 | 1 |
| | N1 | 17(54.8%) | 14 | |
| Lymphatic vessel invasion | ly0 | 8(44.4%) | 10 | 0.39 |
| | ly1/2 | 23(59.0%) | 16 | |
| Venous invasion | v0 | 18(52.9%) | 16 | 1 |
| | v1/2/3 | 13(56.5%) | 10 | |
| Perineural invasion | ne0 | 3(60.0%) | 2 | 1 |
| | ne1/2/3 | 28(53.8%) | 24 | |
| Age | <65 | 9(56.3%) | 7 | 1 |
| | ≧65 | 22(53.7%) | 19 | |
| Sex | Male | 16(53.3%) | 14 | 1 |
| | Female | 15(55.6%) | 12 | |
| Tumor location | Head | 8(42.1%) | 11 | 0.26 |
| | Body and/or Tail | 23(60.5%) | 15 | |

Associations between IQGAP1 expression and the clinicopathologic characteristics were investigated. *P* values were calculated using the Chi-square test. T1, tumor limited to the pancreas, 2 cm or less in greatest dimension. T2, tumor limited to the pancreas, more than 2 cm in greatest dimension. T3, tumor extends beyond the pancreas but without involvement of the celiac axis or the superior mesenteric artery. N0, no regional lymph node metastasis. N1, regional lymph node metastasis. ly0, no lymphatic vessel invasion. ly1, mild lymphatic vessel invasion. ly2, moderate lymphatic vessel invasion. v0, no venous invasion. v1, mild venous invasion. v2, moderate venous invasion. v3, severe venous invasion. ne0, no perineural invasion. ne1, mild perineural invasion. ne2, moderate perineural invasion. ne3, severe perineural invasion.

**Supplementary File 1 Table 4.** Clinicopathological features of high and low ARL4C expression in PDAC with high IQGAP1 expression.

| **Parameters** | | **High ARL4C cases** | **Low ARL4C cases** | ***P* value** |
|---|---|---|---|---|
| Stage | IA-IIA | 11(78.6%) | 3 | 0.3 |
| | IIB-III | 16(94.1%) | 1 | |
| T classification | T1/2 | 3(60.0%) | 2 | 0.11 |
| | T3 | 24(92.3%) | 2 | |
| N classification | N0 | 11(78.6%) | 3 | 0.3 |
| | N1 | 16(94.1%) | 1 | |
| Lymphatic vessel invasion | ly0 | 7(87.5%) | 1 | 1 |
| | ly1/2 | 20(87.0%) | 3 | |
| Venous invasion | v0 | 15(83.3%) | 3 | 0.62 |
| | v1/2/3 | 12(92.3%) | 1 | |
| Perineural invasion | ne0 | 1(33.3%) | 2 | 0.037 |
| | ne1/2/3 | 26(92.9%) | 2 | |
| Age | <65 | 6(66.7%) | 3 | 0.063 |
| | ≧65 | 21(95.5%) | 1 | |
| Sex | Male | 15(93.8%) | 1 | 0.33 |
| | Female | 12(80.0%) | 3 | |
| Tumor location | Head | 20(87.0%) | 3 | 1 |
| | Body and/or Tail | 7(87.5%) | 1 | |

Associations between ARL4C expression in PDAC with high IQGAP1 expression and the clinicopathologic characteristics were investigated. *P* values were calculated using the Chi-square test. T1, tumor limited to the pancreas, 2 cm or less in greatest dimension. T2, tumor limited to the pancreas, more than 2 cm in greatest dimension. T3, tumor extends beyond the pancreas but without involvement of the celiac axis or the superior mesenteric artery. N0, no regional lymph node metastasis. N1, regional lymph node metastasis. ly0, no lymphatic vessel invasion. ly1, mild lymphatic vessel invasion. ly2, moderate lymphatic vessel invasion. v0, no venous invasion. v1, mild venous invasion. v2, moderate venous invasion. v3, severe venous invasion. ne0, no perineural invasion. ne1, mild perineural invasion. ne2, moderate perineural invasion. ne3, severe perineural invasion.

**Supplementary File 1 Table 5.** The sequences of ARL4C ASOs used in this study.

| ASO | Sequence |
| --- | --- |
| Control ASO | T(Y)^a^g^A(Y)^g^a^G(Y)^t^a^5(Y)^c^c^A(Y)^t^c |
| ARL4C ASO-1316 | G(Y)^5(Y)^A(Y)^t^a^c^c^t^c^a^g^g^T(Y)^A(Y)^a |

Lower case=DNA; N(Y)=AmNA; 5(Y)=AmNA_mC; ^=Phosphorothioated

**Supplementary File 1 Table 6.** List of antibodies used in this study.

| | | | Application (Dilution ratio) | | |
|---|---|---|---|---|---|
| Antigen | Company | Catalog # | WB | IHC | ICC |
| ARL4C | Atlas Antibodies (Bromma, Sweden) | #HPA028927 | 1:1000 | 1:50 | |
| Clathrin | BD Biosciences (San Jose, CA, USA) | #610500 | 1:1000 | | |
| EGR1 | Cell Signaling Technology (Beverly, MA, USA) | #4153S | 1:1000 | | |
| β-catenin | BD Biosciences (San Jose, CA, USA) | #610154 | 1:1000 | | |
| Ras (G12D) | Cell Signaling Technology (Beverly, MA, USA) | #14429S | 1:1000 | | |
| Hsp90 | BD Biosciences (San Jose, CA, USA) | #610419 | 1:1000 | | |
| HA | BioLegend (San Diego, CA, USA) | #901502 | 1:1000 | | |
| GFP | Life Technologies/Thermo Fisher Scientific (Carlsbad, CA, USA) | #A6455 | 1:4000 | | |
| GFP | Santa Cruz Santa Cruz Biotechnology (Dallas, TX, USA) | #sc-9996 | 1:1000 | | |
| FLAG | WAKO (Tokyo, Japan) | #014–22383 | 1:1000 | | |
| IQGAP1 | Santa Cruz Santa Cruz Biotechnology (Dallas, TX, USA) | #sc-376021 | 1:1000 | 1:800 | 1:100 |
| MMP14 | Abcam (Cambridge, UK) | #ab51074 | 1:1000 | 1:200 | 1:100 |
| Cytohesin2 | Proteintech Group, Inc (Chicago, IL, USA) | #67185-1-Ig | | | 1:100 |
| Rac1 | BD Biosciences (San Jose, CA, USA) | #610651 | 1:1000 | | |
| Cdc42 | Cell Signaling Technology (Beverly, MA, USA) | #2466S | 1:1000 | | |
| CK19 | Abcam (Cambridge, UK) | #ab52625 | | 1:100 | |
| Mitochondria | Merck Millipore (Billerica, MA, USA) | #MAB1273 | | 1:100 | |
| LYVE-1 | Abcam (Cambridge, UK) | #ab14917 | | 1:100 | |
| YAP/TAZ | Cell Signaling Technology (Beverly, MA, USA) | #8418S | | | 1:100 |

WB, western blotting; IHC, immunohistochemistry; ICC, immunocytochemistry

**Supplementary File 1 Table 7.** Target sequences for siRNA used in this study.

| Gene | Sequence |
|---|---|
| Randomized Control | 5'-CAGTCGCGTTTGCGACTGG-3' |
| human *IQGAP1#1* | 5'-GCTGCACATAGTTGCCTTT-3' |
| human *IQGAP1#2* | 5'-CCCTAATGTAGAATGTCAT-3' |
| human *CYTH2#1* | 5'-GGATGGAGCTGGAGAACAT-3' |
| human *CYTH2#2* | 5'-GCAGTTTCTATGGAGCTTT-3' |
| human *MMP14#1* | 5'-GCAGCCTCTCACTACTCTT-3' |
| human *MMP14#2* | 5'-CCGACATCATGATCTTCTT-3' |
| human *KRAS#1* | 5'-GCATCATGTCCTATAGTTT-3' |
| human *KRAS#2* | 5'-GTTGGAGCTGATGGCGTAG-3' |
| human *CTNNB1#1* | 5'-CCCACTAATGTCCAGCGTT-3' |
| human *CTNNB1#2* | 5'-GCATAACCTTTCCCATCAT-3' |

**Supplementary File 1 Table 8.** Primer sequences for quantitative PCR used in this study.

| Gene | Forward | Reverse |
|---|---|---|
| human *GAPDH* | 5'-TCCTGCACCACCAACTGCTT-3' | 5'-TGGCAGTGATGGCATGGAC-3' |
| human *B2M* | 5'-TGCTGTCTCCATGTTTGATGTATC-3 | 5'-TCTCTGCTCCCCACCTCTAAG-3' |
| human *ARL4C* | 5'-AGGGGCTGTGAAGCTGAGTA-3' | 5'-TTCCAGGCTGAAAAGCAGTT -3' |

*B2M*, *β2-microglobulin*.
